# Rapid assembly of a polar network architecture drives efficient actomyosin contractility

**DOI:** 10.1101/2020.12.08.406298

**Authors:** Vlad Costache, Serena Prigent Garcia, Camille N. Plancke, Jing Li, Simon Begnaud, Shashi Kumar Suman, Anne-Cécile Reymann, Taeyoon Kim, François B. Robin

## Abstract

Actin network architecture and dynamics play a central role in cell contractility and tissue morphogenesis. Pulsed contractions driven by RhoA represent a generic mode of actomyosin contractility, but the mechanisms underlying (1) how their specific architecture emerges, and (2) how this architecture supports the contractile function of the network, remain unclear. Here, we combine quantitative microscopy, single-molecule imaging, numerical simulations and simple mathematical modelling, to explore the dynamic network architecture underlying pulsed contraction. We show that during pulsed contractions, two subpopulations of formins are recruited by RhoA from the cytoplasm and bind to the cell surface in the early *C. elegans* embryo: *recruited* formins, a functionally inactive population, and *elongating* formins, which actively participate in actin filaments elongation. Focusing on formin dynamics during pulses, we show that minority elongating formins precede recruited formins, a kinetic dynamics compatible with formins capturing and rapidly saturating barbed ends available for filament elongation. We then show that these elongating formins assemble a polar network of actin, with barbed ends pointing out of the pulse, pointing to a kinetic rather than mechanical control of network architecture. Finally, our numerical simulations demonstrate that this geometry favors rapid network contraction. Our results thus show that formins saturate available actin filaments barbed ends and convert a local, biochemical gradient of RhoA activity into a polar network architecture, thereby driving rapid and efficient network contractility, an important evolutionary feature in a metazoan with a rapid embryonic cell cycles.

**Highlights:**

1. The formin CYK-1 drives actin network assembly during RhoA-driven pulses
2. The process is extremely rapid, with a formin-based actin elongation rate higher than 1.3 μm·s^-1^
3. A barbed-end saturation mechanism allows for responsive F-actin assembly
4. Rapid and responsive F-actin elongation results in the assembly of aster-like polar actin networks
5. Numerical simulations show network polarity drives very efficient network contractility

## Introduction

Vastly conserved in eukaryotes, the actomyosin cytoskeleton is a major determinant of the mechanical properties of embryonic cells and tissues (Munjal & Lecuit 2014). Modulation of actomyosin networks activity plays a critical role in cell shape changes, cell division, cell migration and polarization. The integration of these behaviors, at the tissue scale, drive tissue deformation and morphogenesis (Lecuit & Lenne 2007). At the molecular scale however, the role of the architecture of actomyosin networks has been a research focus and subject to some debate (Blanchoin et al. 2014; Koenderink & Paluch 2018; Agarwal & Zaidel-Bar 2019). In muscle, the mechanisms for actomyosin contractility has been historically well-characterized, showing that in this quasi-crystalline organization, the sliding of bipolar Myosin II mini-filaments along actin filaments drives network contractility. In other cell types however, and in particular in the cell cortex of developing embryos, the seemingly disordered actin network remains poorly understood in terms of network polarity, length distribution, mesh size, turnover rates or crosslinking levels, and we still do not fully understand how F-actin architecture is linked to network contractility. Theoretical studies (Galkin et al. 2010; Galkin et al. 2011; Lenz, Gardel, et al. 2012) and computational models (T. Kim 2015) have shown that asymmetry between compressive and extensive modulus—the ability to withstand tension but buckle under compressive forces—can drive contraction of disordered bundles. Similarly, numerical simulations and *in vitro* experiments have clearly demonstrated that non-polar actin networks can contract (Yu et al. 2018). Cellular networks however often display characteristic organizations, suggesting that specific network dynamics and geometries may play a critical role in network contractility (Koenderink & Paluch 2018).

RhoGTPase zones have recently emerged as essential regulators to template the architecture of the actomyosin meshwork by defining active, task-tuned zones of cytoskeletal assembly (Benink 2005; Miller & Bement 2009; Burkel et al. 2012; Bement et al. 2005). Examples of such zones include the leading edge of migrating cells, the cleavage furrow during cell division, or the apical cortex during apical constriction. During embryonic morphogenesis in particular, a wide class of morphogenetic processes are driven by brief iterative contractions of the cortical actomyosin network termed pulsed contractions (He et al. 2010; H. Y. Kim & Davidson 2011; Martin et al. 2009; Rauzi et al. 2010; Munro et al. 2004; Roh-Johnson et al. 2012). Previous work showed that pulsed contractions are driven by excitable dynamics of the Rho GTPase RhoA, leading to the formation of activation zones that drive the recruitment of downstream effectors formin, Anillin, F-actin and Myosin II (Maddox et al. 2005; Munro et al. 2004; Michaux et al. 2018; Naganathan et al. 2018; Reymann et al. 2016). Excitable dynamics, with a feedforward activation and delayed negative feedback, seem to play an important role to establish Rho activation (Bement et al. 2015; Maître et al. 2015; Nishikawa et al. 2017; Michaux et al. 2018).

RhoGTPases thus spatially and temporally pattern the recruitment, turnover and activity of downstream effectors. It remains unclear, however, how these orchestrated modulations of actomyosin dynamics support the specific cellular function of Rho zones. Here, we show that the dynamics and topology of RhoA activation, converting a RhoA chemical gradient into the assembly of a polar actin network, drives the formation of a network structure tuned to its contractile function.

In the nematode *C. elegans*, pulsed contractions occur from the 1-cell stage onwards during interphase (Munro et al. 2004; Mayer et al. 2010) and support cell polarization and apical constriction (Nance & Priess 2002; Nance 2003; Roh-Johnson et al. 2012). Here, we show that RhoA pulses control the accumulation of the formin CYK-1 (diaphanous/mDia homolog), driving F-actin accumulation during pulsed contractions. We further show that actin network assembly is kinetically controlled by the saturation of actin filaments barbed ends, resulting in a time-optimal response to RhoA activation. Using single-molecule microscopy to infer local actin filament orientation during pulse assembly, we show that formin-assembled actin networks are polar, generating networks with barbed ends pointing outside of the pulse. Finally, our computational exploration shows that this polar network architecture is favorable to the generation of efficient actomyosin contractility. Taken together, these results underline a kinetic rather than mechanical control for actomyosin network orientation during pulsed contractions. They also underline the tinkering evolution of billion-years old machinery, reusing the molecular machines—formin, F-actin and Myosin II—to drive a fundamentally conserved phenomenon, precisely-tuned force generation, with opposite geometries reflecting organism-specific construction rules and constraints.

## Results

### Cortical dynamics of formins in a developing embryo

Formins are actin nucleators and processive actin elongators, catalyzing the addition of actin monomers to the barbed end of actin filaments while protecting the filament against capping (Pruyne et al. 2002). In *C. elegans*, 7 formin genes have been identified (Mi-Mi et al. 2012). Among these, *cyk-1* (cytokinetic defective-1), the only ortholog of the Diaphanous family of formins, is required for cell division (Swan et al. 1998).

To study CYK-1 in the early *C. elegans* embryos, we first used CRISPR-Cas9 homologous recombination to insert a GFP in the genomic *cyk-1* locus. We then used live single-molecule fluorescence microscopy to visualize the dynamics of individual formin molecules fused with GFP (Robin et al. 2014). We first observed that formins apparently classified in at least two populations (Movie S1), a population of ballistic molecules and a static population. To better visualize these two populations, we used maximum intensity projection to overlay the position of molecules over 100 consecutive time-points (Movie S2). Using this visualization tool, static formins appeared as dots, while moving formins appeared as a trail on the cell surface.

To quantitatively characterize these two populations, we performed single-particle tracking and analyzed the trajectories of 19137 individual formin molecules from 5 embryos. Based on the logarithmic regression of the mean-squared displacement to an anomalous diffusion model *MSD = 2*D.t^α^* (Robin et al. 2014), we characterized all particle trajectories longer than 15 frames (Fig. 1A,B) by their anomalous diffusion coefficient *D* and scaling exponent *α*. Strikingly, we observed the emergence of two clear populations, corresponding to the static and ballistic populations, with apparent distributions of scaling exponent peaking at α = 0.3 (subdiffusive) and α = 1.6 (superdiffusive), respectively (Fig. 1C,D, Movie S3).

**Figure 1.**
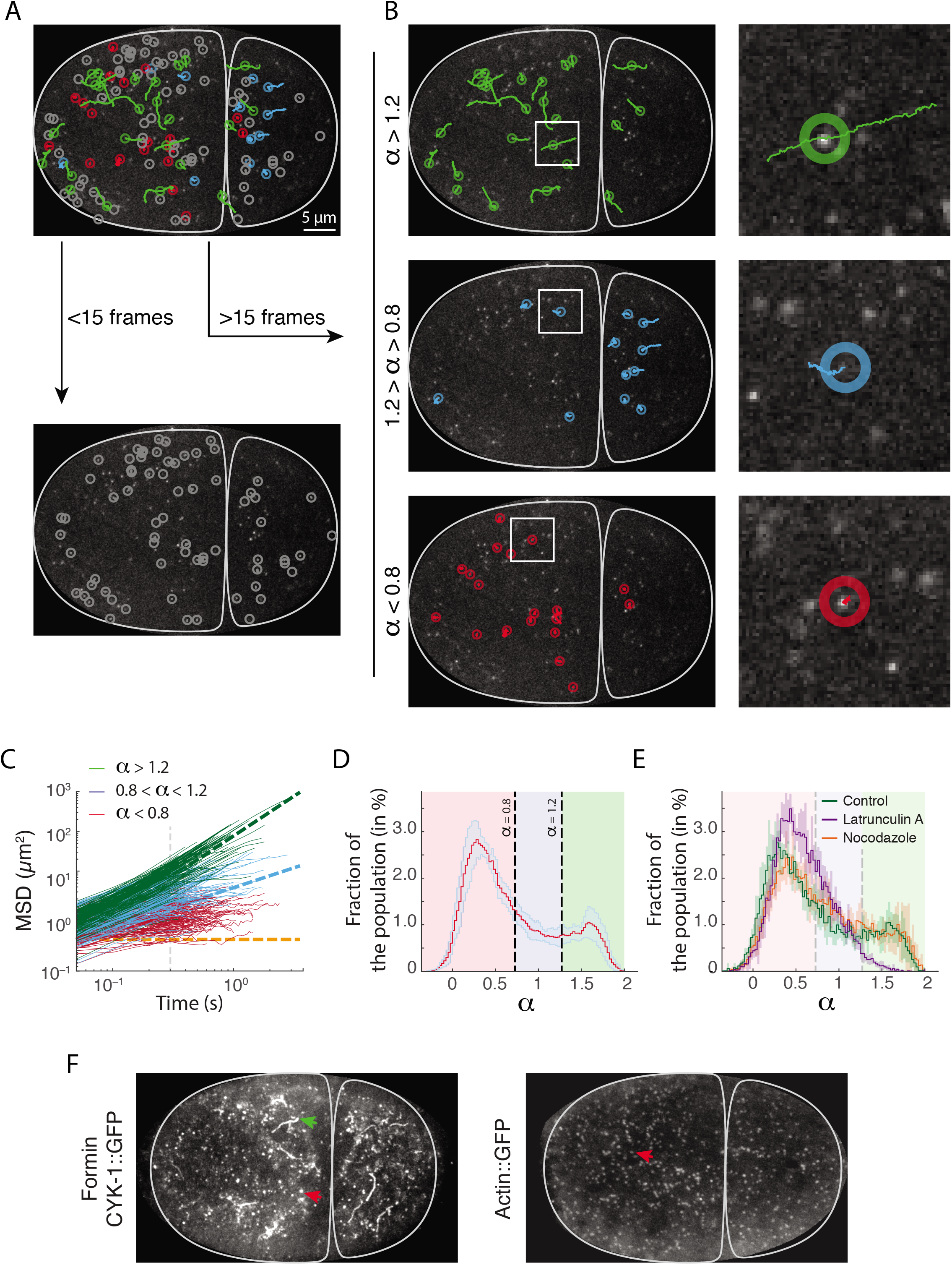
Anomalous diffusion of individual formin molecules identifies a subpopulation of actin-elongating formins. **(A)** Single-molecule imaging and tracking of formins fused with GFP (CYK-1::GFP) shows individual behaviors ranging from superdiffusive (green) to diffusive (blue) to subdiffusive (red). **(B)** Mean-square displacement against lag time. Slope curve reports on the anomalous diffusion exponent. Particles with anomalous diffusion exponent larger than 1.2 in green, between 0.8 and 1.2 in blue, and smaller than 0.8 in red. Pure superdiffusive corresponds to α = 2 (green dashed), pure diffusive α = 1 (blue dashed), and immobile (orange dashed). **(C)** Distribution of the fraction of particles displaying a given anomalous diffusion exponent in 5 movies (average +/- SD). Background shows the domains corresponding to the classification used here. Two peaks seem to emerge, centered at α = 0.3 and α = 1.6. **(D)** Detected mobilities correspond to different classes of behaviors. Superdiffusive display a characteristic ballistic motion (green, top panel), while subdiffusive particles appear immobile in the cortex (red, bottom). **(E)** Compared to control (green curve), the superdiffusive population is absent in embryos treated with Latrunculin A (purple), but not Nocodazole (orange). More than 2000 tracks analyzed per embryo for, with >5 embryos per condition presented. **(F)** Projection over 5s (100 consecutive frames) of formin CYK-1::GFP (left) and Actin::GFP (right) speckles, showing subdiffusive speckles (red arrow) and superdiffusive trails (green arrow). Actin::GFP does not display superdiffusive trails.

Previous work suggested that these super-diffusive particles represented formins actively elongating actin filaments (Higashida 2004; Funk et al. 2019). To confirm this, we used RNAi against *perm-1*, a known component eggshell protein (Carvalho et al. 2011; Olson et al. 2012), to permeabilize the eggshell, and subsequently treated the embryos with the microtubule depolymerizing drug Nocodazole and the actin depolymerizing drug Latrunculin A (Movie S4). Performing the same analysis as previously, we observed that the superdiffusive population essentially disappeared after Latrunculin A treatment, while it was unaffected by Nocodazole treatment (Fig. 1E, Movie S4). These results strongly supported the idea that superdiffusive cortical CYK-1::GFP speckles corresponded to formin dimers actively and processively elongating actin filaments at the barbed-end of the filament at the cell cortex.

To measure the speed of formins, we selected a collection of trajectories projected formin motion on a smoothed version of their trajectory, and quantified the traveled distance along this trajectory, or using MSD measurements presented before. Both metrics, quite conservative, yielded very similar result of 1.1 ± 0.2 μm·s^-1^ and 1.3 ± 0.2 μm·s^-1^ (standard deviation)—in line with previously reported speeds (Higashida 2004), but slower than recent *in vivo* reports (Funk et al. 2019). Interestingly, single-molecule microscopy of actin::GFP speckles, serving as fiducial markers on the network of actin filaments, remained largely immobile (Fig. 1F, Movie S5, (Robin et al. 2014)), supporting the idea that actin filaments are not extruded by immobile formins, and that filament elongation instead fully translates in formin directional motion. These data show that CYK-1 velocity is a reliable *in vivo* proxy for formins elongation rate, demonstrating an average elongation rate of ~400–468 monomers·s^-1^. Incidentally, our results also suggest that CYK-1 could be used as a biosensor to measure cellular modulations of the concentrations of profilin-ATP-G-Actin, calibrated on elongation rates previously reported *in vitro* in the presence of profilin (Neidt, Scott, et al. 2008; Neidt, Skau, et al. 2008). Provided that in our system formin-mediated actin filament elongation rates are not buffered by slow dissociation of profilin from the barbed end (Funk et al. 2019), or modulated by mechanical forces (Jégou et al. 2013; Courtemanche et al. 2013; Kubota et al. 2017), our results would point to a local G-actin concentration in the early embryo in the ~10-12 μM range.

### Formin-mediated actin filament elongation rates during the cell cycle

To explore if actin elongation was dynamically modulated during embryonic development, we then measured formin velocity at the 1-, 2- and 4-cell stages during distinct phases of the cell cycle (Fig. 2A, see Material and Methods). To avoid confounding effects of overcrowding on tracking at high particle density, we decided to use a strain over-expressing GFP fused with CYK-1, which displays essentially identical dynamics as GFP fusion at the endogenous site, but allowed us to visualize formins at much lower densities, improving particle tracking. At the 1-cell stage, elongation rate remained unchanged from polarization to maintenance phase (1.25 ± 0.18 μm·s^-1^ and 1.27 ± 0.16 μm·s^-1^, resp.), but decreased significantly over the entire cortex during cytokinesis, to 1.05 ± 0.22 μm·s^-1^.

**Figure 2.**
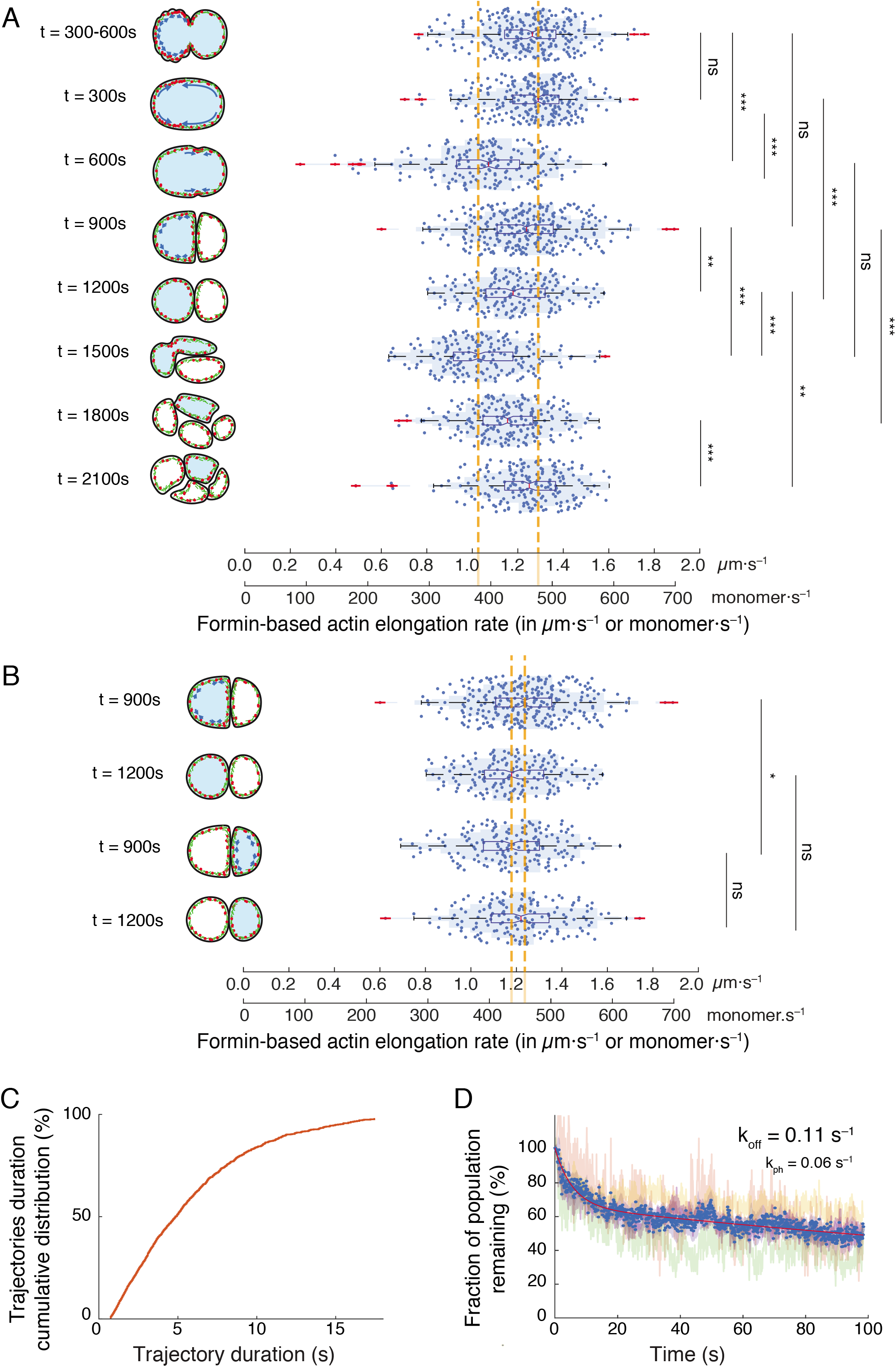
Formin speed is changed by the cell cycle but is conserved through cell lineage. **(A-B)** Formin speed in AB (A) and P1 (B) cell. Right: Distribution of elongating Formins speed. Left: Schematic of the stage and location of the cell from which the tracks are extracted, with measured cell in light blue. Myosin in red, Actin in green. * ≥ 0.05, ** ≥ 0.01, *** ≥ 0.001, ns: not significant. Dashed orange line marks upper and lower averages. Outliers in red. (See Table S1,2 for detailed statistical information). **(C)** Cumulative distribution of CYK-1::GFP trajectory duration, as a fraction of detection events, showing a half-life of ~5 s. **(D)** The surviving fraction of cortical CYK-1::GFP as a function of photobleaching time reports both on the turnover rate koff and the photobleaching rate kph. Bi-exponential fit in solid red. Experiments performed on strain over-expressing CYK-1 fused with GFP.

The velocity increased again after cytokinesis, going back to its level at corresponding 1 −cell stage polarization establishment (1.24 ± 0.19 μm·s^-1^), stable or slightly decreasing during mitosis (1.18 ± 0.18 μm·s^-1^) in the AB cell, to drop down again during AB cytokinesis (1.04 ± 0.18 μm·s^-1^). Again, during interphase at the 4-cell stage in ABp, formin velocity increased again though notably lower in interphase compared to mitosis (1.15 ± 0.17 μm·s^-1^ at interphase and 1.24 ± 0.18 μm·s^-1^ during mitosis). Taken together, these results show that formin speed drops significantly during cytokinesis but increases when interphase resumes.

To compare actin elongation rates across lineages, we also measured formin speed in AB and P1 at the 2-cell stage during interphase (1.24 ± 0.19 μm·s^-1^ and 1.18 ± 0.18 μm·s^-1^, respectively), and during mitosis (1.18 ± 0.18 μm·s^-1^ and 1.22 ± 0.19 μm·s^-1^, resp.) (Fig. 2B). Our results suggest that formin speed does not vary between the two cell types.

In summary, actin elongation dynamics is distinctly modulated during phases of the cell cycle, decreasing significantly by >10% during cytokinesis, to increase again after cytokinesis completion. Strikingly, measured elongation rates seemed relatively robust and only changed marginally from the 1-cell to 4-cell stage. These results suggest that F-actin dynamics might be differentially regulated by G-actin concentration, but remains largely robust across cell lineages during early embryonic development.

### Formin kinetics and implications on actin filament length *in vivo*

The length of formin-elongated actin filaments is controlled by a combination of the formin elongation rate, formin global off-rate (combining unbinding and competition), and the turnover rate of F-actin monomers in the cortex. Specifically, we assumed that F-actin turnover and elongation dynamics are independent processes and follow exponential laws with characteristic rate 1/τ_actin_ and 1/τ_formin_. For an F-actin monomer at the time of its disassembly, the length of filament from monomer the to the barbed-end of the filament follows an exponential law with characteristic length L_filament_ = V_formin_ × τ_filament_, where 1/τ_filament_ 1/τ_actin_ + 1/τ_formin_ is a characteristic “off” rate of formins and V_formin_ their speed (see Suppl. Material for detailed derivation). To estimate filament length, we thus needed to access actin and formin off-rates, and use our measured actin filament elongation rates.

We expected a significant fraction of trajectory to be interrupted, either by tracking failure or photobleaching, barring us from using single-molecule tracking as a proxy (Fig. 2C). We therefore turned to a previously established strategy, smPReSS (Robin et al. 2014) to estimate a bulk turnover rate for formins. Briefly, by measuring the depletion of cortical formins caused by laser illumination of the cortex in a CYK-1 overexpression strain, we can estimate a *bulk* formin turnover rate, over all formin populations. We could thus establish that the bulk cortical turnover rate of the formin CYK-1 is ~0.11 s^-1^ (Fig. 2D).

Combining our results with previous measurements of Actin::GFP turnover rates (0.05–0.15 s^-1^, (Robin et al. 2014; Michaux et al. 2018)), we estimate that formin-elongated actin filaments scale to ~6 μm on average in the 2-cell stage *C. elegans* embryo.

### Dynamics of formin at the cortex during pulsed contractions

We then decided to focus on the dynamics of formins during pulsed contractions. During polarity establishment in the 1-cell embryo, and during interphase at the 2-cell stage, formins accumulate in well-identifiable pulses corresponding to RhoA-driven actomyosin pulsed contractions (Fig. 3A-E, Movie S2, Fig. S1A, (Munro et al. 2004; Piekny et al. 2005; Naganathan et al. 2014; Reymann et al. 2016; Michaux et al. 2018)). To infer the biochemical sequence of formin activation during pulsed contractions, we thus decided to measure the timing of arrival of the various formin populations over the course of a pulsed contraction.

**Figure 3.**
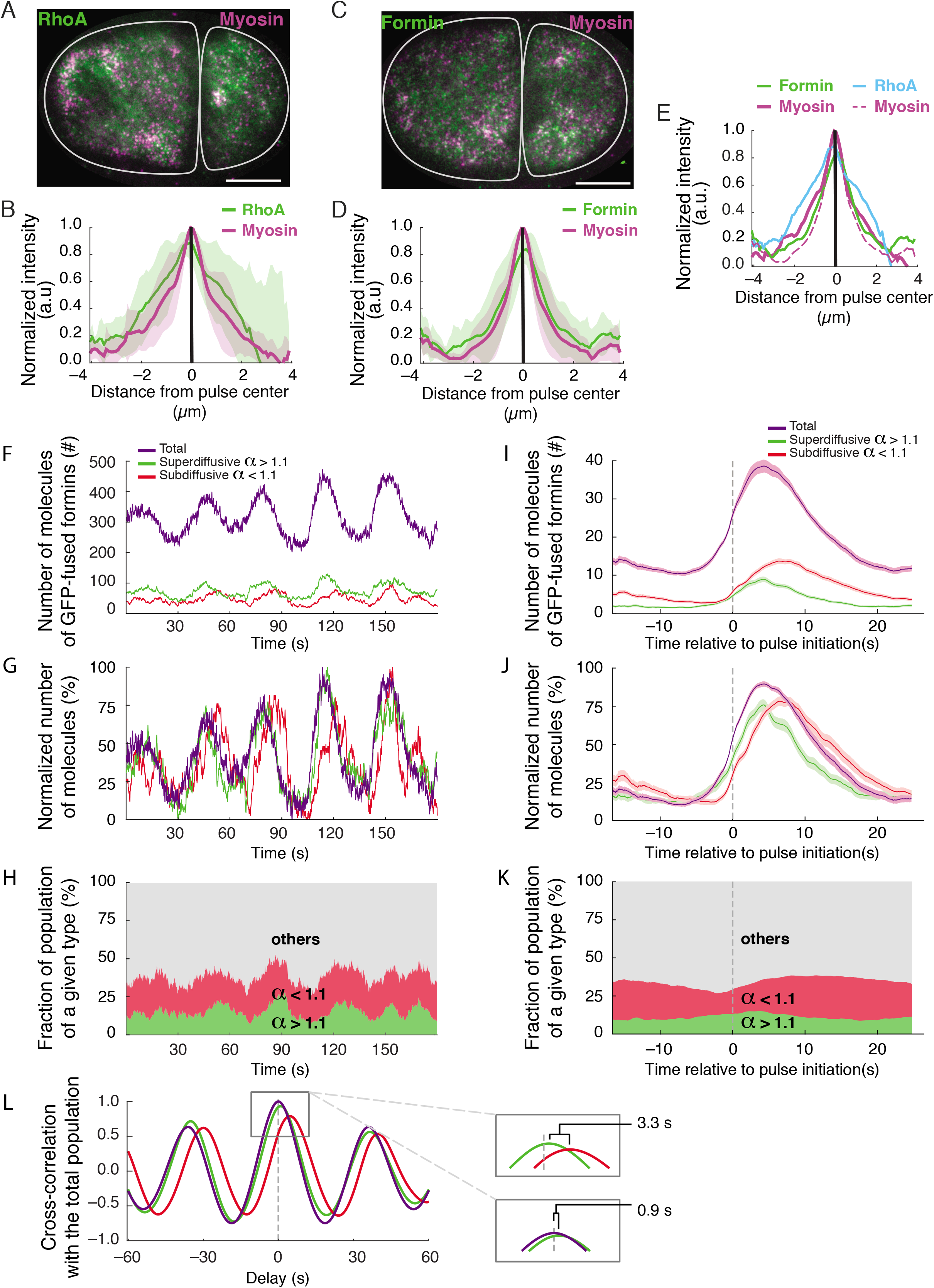
Recruited and actin-filament elongating formins display distinct dynamics during pulsed contractions. **(A)** Two-cell stage embryo showing RhoA biosensor (green, AHPH::GFP) and Myosin II (magenta, NMY-2::mKate2 at endogenous locus). **(B)** Mean normalized intensity of RhoA and myosin density profile across pulses. **(C)** Two-cell stage *C. elegans* embryo expressing fluorescent protein fusions of formin fused with GFP in green (CYK-1::GFP) and myosin in magenta (NMY-2::mKate2), at the endogenous loci. **(D)** Formin normalized mean intensity (solid green) and myosin (solid magenta) density profile along an axis drawn through pulses. **(E)** Compiled results from (B,D). (A,C) scale bar: 10 μm. (B,D,E) shaded curves represent standard deviations from 18 pulses from 6 embryos. **(F-H,L)** Population dynamics of formin CYK-1 fused with GFP, in a single embryo during 5 consecutive pulsed contractions. **(F)** Number of total (purple), super-diffusive (green) and subdiffusive (red) GFP-fused formins molecules during pulsed contractions varies in a periodic manner. **(G)** Normalized number of molecules during pulsed contractions. The populations display distinct accumulation dynamics. **(H)** Temporal evolution of the relative fraction of superdiffusive (green) and subdiffusive (red) subpopulations within the total population during pulsed contractions. **(I-K)** Dynamics of formin populations during pulsed contractions averaged over 115 pulses from 10 embryos. Individual pulses are synchronized to pulse initiation (t=0, see also Fig. 5F). **(I)** Number of total (purple), superdiffusive (green) and subdiffusive (red) GFP-fused formins molecules during pulsed contractions. **(J,K)** Absolute (J) and normalized (K) number of molecules during pulsed contractions shows that superdiffusive (green), subdiffusive (red) and total formin (purple) populations accumulate with distinct dynamics. **(L)** Cross-correlation with total population of superdiffusive (green), subdiffusive (red), and total (purple) formin populations. Offset show that subdiffusive formins accumulate 3.3 s after superdiffusive. (F-L) Strain over-expressing CYK-1 fused with GFP.

As described previously, in order to categorize into subdiffusive or superdiffusive, a minimal track length was required. We thus divided the population into 3 tiers: short tracks (<15 consecutive time frames), which could not be categorized into a specific population, long subdiffusive and long superdiffusive. Using this technique, we were able to demonstrate that the ratio between the different populations was finely modulated during pulses (Fig. 3F-K). To characterize the dynamics of arrival of these populations at the cell cortex, we first focused on the kinetics of these populations on a sequence of successive pulses (Fig. 3F). Strikingly, we observed an iterated sequence of accumulation (Fig. 3G,H). Using cross-correlation, we measured a delay between the arrival of the superdiffusive and subdiffusive populations of ~3 s (Fig. 3L). This suggested that the distinct populations accumulated at the cortex in a sequence, superdiffusive formins (hereon, *elongating* formins) accumulating first, followed by subdiffusive formins (hereon *recruited* formins).

To confirm this result, we collected a series of 115 pulses from 10 embryos, and quantified the dynamics of the different formin subpopulations. Based on these results, we observed that formins indeed accumulated at the cortex in a well-defined sequence, starting with superdiffusive followed by subdiffusive formins (Fig. 3I-K, Movie S6). We further confirmed this observation using a different metrics (based directly on particle displacements instead of trajectory classification) to measure this delay (Fig. S1B-E), and yielding very similar delays (Fig. S1F-I).

This result was somewhat surprising, as based on previous work on formin structure and domain activity, we expected an activation sequence whereby formins would be first recruited to the cortex by RhoA, then transferred to barbed-ends of actin filaments to promote elongation (F. Li & Higgs 2005; Higgs 2005; F. Li & Higgs 2003). Numerically however, the number of recruited formins out-weighted the elongating population (see Fig. S1F, Class 1 vs. Class 2), suggesting that the system might be running in a regime in which formins are in excess, and elongate a limiting pool of barbed ends available for elongation.

### A barbed end saturation mechanism allows for responsive actin assembly

To test this hypothesis, we designed a simple kinetic model for CYK-1 recruitment, and used this model to explore the temporal dynamics of formin accumulation (Fig. S2A,B). We postulated that:

(1) active RhoA concentration pulses periodically, with period 30 s (Michaux et al. 2018),
(2) cytoplasmic formins are activated by active RhoA and *recruited* to the cortex, shifting in the “**recruited”** population (F. Li & Higgs 2005),
(3) CYK-1 formins are poor nucleators but good elongators—we considered that formins do not efficiently nucleate new filaments under physiological conditions (*in vitro* actin assembly yields ~1 new nucleated filament per 550 CYK-1 formin molecule at 2.5 μM actin and 2.5 μM profilin PFN-1 (Neidt, Skau, et al. 2008)),
(4) once recruited at the cortex, formins bind to barbed ends through a bimolecular reaction to drive actin assembly, becoming **“elongating”** formins,
(5,6) recruited and elongating formins unbind from the cortex, returning back to the cytoplasmic pool with characteristic rates 1/τ_recruited_ and 1/τ_elongating_.

To seed our model, we used measured parameter values for RhoA activity, formin unbinding rates and relative ratios between the different formin populations. Using these parameters, and provided that in our parameters (1) the binding reaction of recruited formins to barbed ends is very fast, and (2) barbed ends are scarce and are depleted when formin density increases, our model indeed captured the key observation that elongating formins accumulated before recruited formins (Fig. 4A-C). Indeed, under these conditions, during an early phase elongating formins accumulate rapidly following the RhoA pulse, followed by a late phase during which recruited formins accumulate (Fig. 4B,F, Fig. S2C-H).

**Figure 4.**
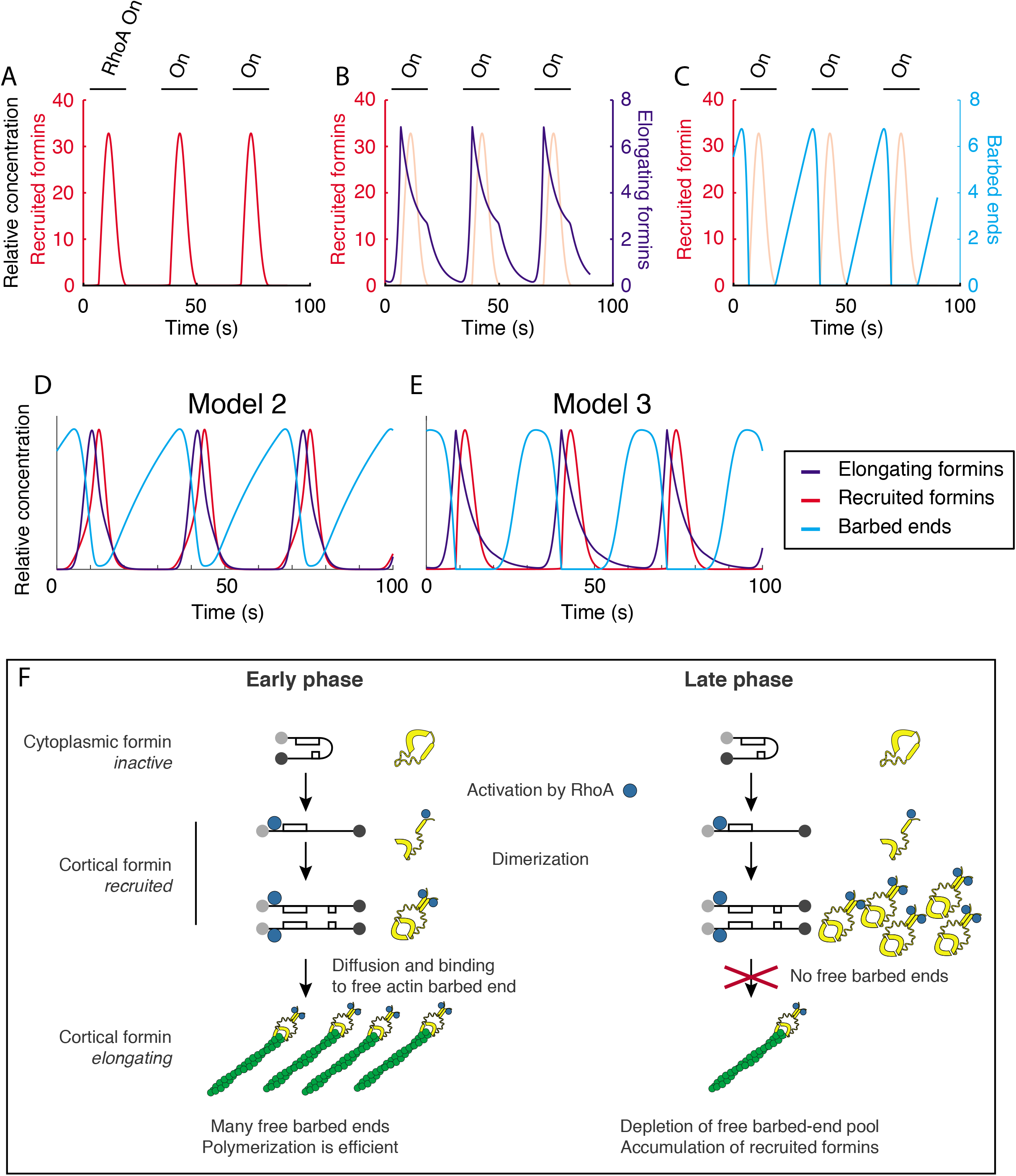
Modeling formin recruitment dynamics with an excess of formins over barbed ends reproduces *in vivo* recruitment sequence. **(A)** Temporal dynamics of recruited formins during a sequence of 3 pulses. Red: Recruited formins. **(B)** Same, with elongating formins (purple: elongating formins, light red: recruited formins). Recruited formins accumulate after elongating formins. **(C)** Temporal dynamics of barbed ends (Cyan: barbed ends, light red: recruited formins). The model uses two free parameters and 4 parameters set based on experimental measurements. Barbed ends accumulated progressively in the absence of formins (between pulses), but are rapidly used upon formins recruitment. Recruited formins are immediately converted in elongating formins, such that elongating formins accumulate until depletion of the built-up barbed ends (C, purple). RhoA activation period is denoted by a black line and denoted by “Rho On”, or “On”. **(D)** Same model with a different choice of parameter values for the free parameters of the model. Outcome is similar. **(E)** Distinct model, where barbed ends are generated periodically during the end of the pulsed contractions, mimicking a myosin-driven actin buckling/severing activity. This model also readily reproduced the expected outcome without additional refinement. **(F)** Schematic representation of the two phases of the pulse, representing a first-come first served scenario. **Early phase:** formins arrive at the cell surface, barbed ends are available, and recruited formins are immediately converted into elongating formins. **Late phase:** Upon depletion of the barbed end pool, formins are trapped in the recruited state.

We favored a model for activation in which RhoA binding preceded dimerization (Fig. S2A), though other models—e.g. dimer exists before the formin binds to RhoA and unfolds—are also plausible. The simulation however proved robust to these modifications of the biochemical scheme (Fig. 4D,E).

Another class of model could invoke the delayed activation by RhoA of a formin competitor for barbed-end binding. Recently, the capping protein CapZ/CAP-1 was described as forming a *ménage-à-trois* with formins at the barbed, weakening formin-barbed ends binding affinity and eventually leading to formin displacement (Shekhar et al. 2015). While, in this scenario, the shift from elongating to recruited/inactive formins would result from a mechanism relying on competition for barbed ends rather than saturation of barbed ends, such a model would essentially present the same kinetic signature, with a “pulse” of barbed-ends available for polymerization.

These results show that given a small set of assumptions, we could explain the emergence of a significant delay between recruited and elongating formins. This model suggests that the saturation by CYK-1 of the barbed-ends of actin filament allows for a rapid response to pulsed RhoA activation (Fig. S2C-H). This suggests that the kinetics of the actin cytoskeleton in the early *C. elegans* embryo is wired to drive fast response to an upstream activation of actin dynamics.

### Relative rates of actin assembly and contractility support polar network assembly

During pulsed contractions, cortical contractile dynamics results in peak cortical flow rates of ~0.3 μm·s^-1^ (Michaux et al. 2018; Munro et al. 2004; Nishikawa et al. 2017). In comparison, elongating formins move relatively rapidly, with a measured speed of 1.1-1.3 μm·s^-1^. As a consequence, even at the peak of contraction, elongating formins can “exit” the contraction zone easily and assemble an actin network of filaments up to several microns around the pulse region. To describe the architecture of this network, we measured the orientation of formin-based actin elongation during pulse assembly. To this end, we focused on elongating formins, and measured the orientation of elongation radially away from the zone of formin accumulation (Fig. 5A-D), which essentially corresponds to the RhoA recruitment zone (Fig. S1A). Displaying only orientations where we could collect >200 individual elongation measurements, we observed that while formins are not heavily oriented outside of the pulse time-window (Fig. 5E,F). In contrast, during the peak of assembly (approx. corresponding to the period where elongating formins >50% max.), formins displayed a strong polarization (Fig. 5E).

**Figure 5.**
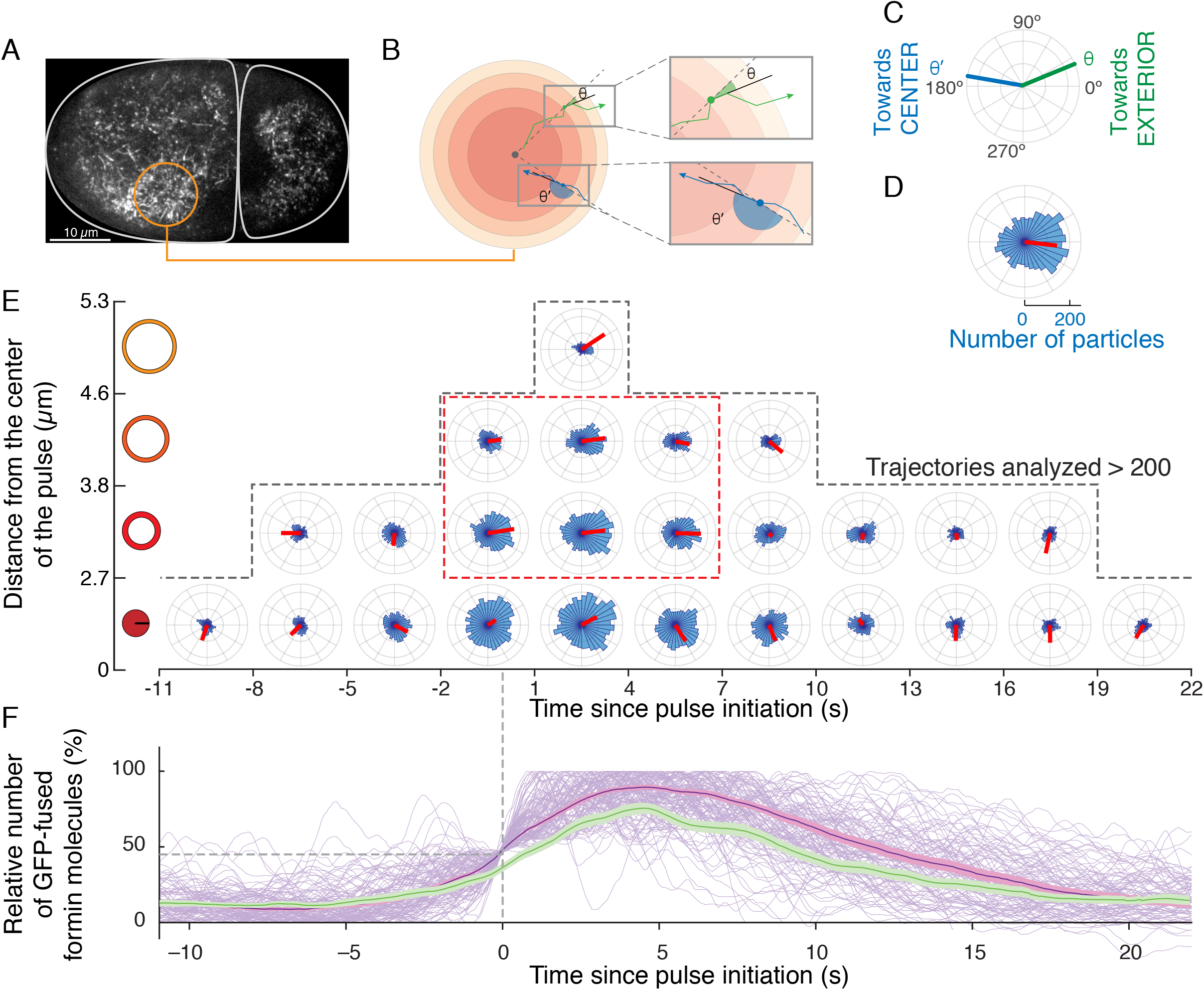
CYK-1 formin-driven actin filament elongation during pulsed contractions drives the formation of a transient polar actin network, barbed ends pointing out. **(A)** Image of a 2-cell stage embryo labelled with CYK-1::GFP and displaying the area corresponding to a pulsed contraction (white dashed line). **(B)** Measure of the angle is performed with respect to the center of the pulse and the local orientation of the formin trajectory. **(C)** Green (resp. blue) track oriented with filament barbed-end pointing away (resp. towards) from the center of the pulse. **(D)** Angle distribution for steps of elongating formins. Average elongation orientation as a red segment. **(E)** Angle distribution of superdiffusive formin trajectory steps during pulsed contractions. **During the peak of the pulse, around pulse center, superdiffusive formins display on average an outwards orientation (dashed orange box).** 115 pulses derived from 10 distinct embryos were used to collect >50 000 trajectories. Steps are binned according to the distance from the center of the pulse (vertical) and time from t=0 (horizontal, 3s intervals) to produce each rose plot. Steps are then mapped in the polar histogram as in (C). Individual pulses are synchronized to pulse initiation (t=0, first pass at 45% of the normalized number of particles in the considered pulse), as shown in (F). Average step orientation displayed as a red segment, length reflecting statistical significance. Rose plots with less than 200 steps not represented (dash line outlines plots with >200 steps). **(F)** Evolution of the number of particles in a pulse. Same axis as (E). In light purple, total number of formin particles in individual pulses. In dark purple (resp.), the corresponding average and SEM. In green, average and SEM for the super-diffusive population. Same dataset as Fig. 3I-K.

These results show that formins elongate the actin network with a polar dynamics, elongation during the pulse occurring from the center of the pulse to the outside. As formin-based elongation increases local actin concentration ~2-fold (Michaux et al. 2018), we propose that pulses assemble a polar actin network with barbed-ends pointing outwards of the pulse akin to an “actin aster” (see discussion).

To test if this orientation resulted purely from the transient local gradient of elongating formins between the pulsing region and its surroundings, or if additional mechanisms should be invoked, we designed a simple spatial model of formin orientation. To seed our model, we exclusively used measured parameters of formin recruitment and elongation dynamics (formin-mediated actin filament elongation rate, density, activation/elongation duration, and off-rate, and pulsed contractions localizations), and generated synthetic formin pulsed accumulations with random orientations. Modelled formin dynamics displayed similar orientations, with filaments pointing outwards, and closely mirroring the dynamics observed *in vivo* (Fig. S3, Movie S7). Altogether, these results demonstrate that local formin accumulation drives the assembly of a polar actin network architecture with a majority of barbed ends pointing out.

### Actomyosin network polarity supports efficient contractility

While previous work, both theoretical (Lenz, Thoresen, et al. 2012; Lenz, Gardel, et al. 2012) and *in vitro* (Linsmeier et al. 2016), showed that actin contraction does not require a specific network orientation, *in vivo* observations suggested that pulsed contractions form a polar actin network (Coravos & Martin 2016). Strikingly, recent *in vitro* and computational work show that Myosin II contractility can drive polar network reorganization by barbed end filament sorting, with an opposite polarity (Kreten et al. 2018; Wollrab et al. 2018). We thus wondered if the polar network architecture we observed, barbed end pointing out— combined with Myosin II intrinsic polarity as a plus-end directed motor (Howard 2001)—would not support either stronger contractions or contraction over larger distances. Controlling independently network orientation and density, while faithfully constraining other parameters, however, was not experimentally manageable *in vivo* or *in vitro*. We decided to turn to agent-based models of cortical mechanics to decipher the impact of network architecture on contractility.

Using our established computational model of the actomyosin networks (Fig. S4A, (Jung et al. 2015; Bidone et al. 2017; T. Kim 2015)), we probed the roles of formin-induced F-actin elongation in cortex mechanics and architecture. Using a cortex-like actin meshwork (20 μm × 20 μm × 100 nm), we simulated RhoA-driven pulsed contraction by locally modulating the kinetics of Myosin II and F-actin elongation rates, based on experimental measurements (Fig. S4B). Specifically, to reproduce formin activity, we increased the elongation rate of a fraction of the barbed ends in the RhoA-activated region, resulting in rapid elongation of actin filaments for ~10 s, or ~12 μm (Fig. 6A, top row). We then locally turned on Myosin II activity in the RhoA-activated region for 15 s, and with a delay of ~5s to reproduce delayed Myosin II activation by RhoA (Movie S8-10, (Michaux et al. 2018)).

**Figure 6.**
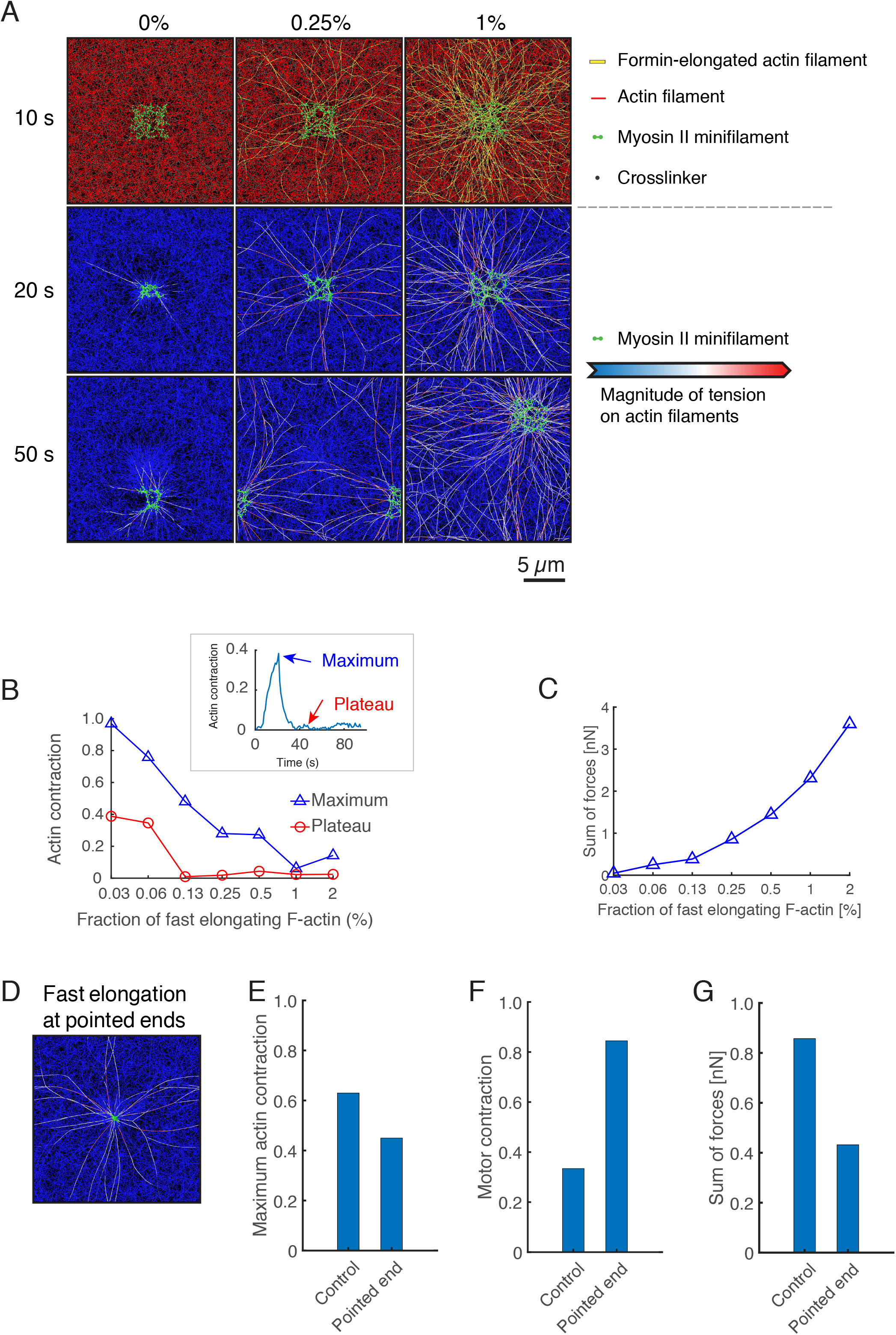
Agent-based numerical simulations of the mechanics of pulsed contractions demonstrate that polar network architecture supports efficient actomyosin contractility. **(A)** Simulating a sequence of two successive pulsed contractions. First pulse occurs in the center, while second pulse location is stochastic. Snapshots taken at *t* = 10, 20, and 50 s (resp. during pulse initiation, first pulse, and second pulse, see Fig. S4B) with three different fractions of barbed ends undergoing quick elongation: 0%, 0.25%, and 1%. Top row: Actin, myosin, and actin cross-linking protein resp. in red, green, and gray. Formin-elongated actin filaments assembled in yellow. Bottom rows: Magnitude of tensile forces on filaments. Green overlay represents active myosin motors. Simulations have periodic boundaries conditions. **(B)** Maximum and plateau values (blue triangles, red circles, resp.; see inset for definition) of the actin contraction as a function of the fraction of fast elongating actin filaments. Contraction is computed in the pulsing region (see Methods for details). Inset shows time evolution of actin contraction (fast elongation at 0.25%), maximum and plateau. **(C)** Sum of tensile forces acting on quickly elongated filaments depending on the fraction of long filaments. **(D)** Snapshot at *t* = 20 s of a network with actin filaments elongated from pointed ends. Color schemes identical to second and third rows of (A). **(E, F, G)** Maximum actin contraction (E), motor contraction (F) and sum of tensile forces on elongated filaments (G) quantified at *t* = 20 s. In the control case, filaments are elongated from barbed ends, whereas in the other case, filaments are elongated from pointed ends.

Using this tailored model of pulsed contraction, we then evaluated the impact of formin activation levels on network architecture and the deployment of forces generated by Myosin II. We observed that actin and myosin tended to contract toward the center of the activated region upon motor activation, peak, then relax towards a plateau upon Myosin II inactivation (Fig. 6B inset, Fig. S4C). Interestingly, the maximum levels of actomyosin contraction decreased with formin activation level (Fig. 6B, Fig. S4D). Meanwhile the sum of forces experienced by formin-elongated actin filaments increased with formin activation level (Fig. 6A(second row),C): long, formin-elongated actin filaments are cross-linked with many other short actin filaments, propagating the force generated by Myosin II farther in the network (Fig. S4E). We also observed that weaker local contraction and long-ranged force transmission in the network prevented the formation of contraction-induced actin aggregates separated from the rest of the network (Fig. 6A-B). At high formin activation levels, myosin and actin contraction were both inhibited, preventing the appearance of aggregates (Fig. 6B, Movie S10). In summary, as more actin filaments are elongated by formin, resistance to contraction increased, preventing local network collapse, while enabling force transmission farther in the cortex.

Finally, in order to explore the specific impact on contractility of network polarity (Fig. 5), we decided to probe the mechanics of network displaying inverted architectures. We therefore set out the numerical simulations to assemble actin networks with pointed ends emerging from the aster (by inverting formin polymerization dynamics, enhancing formin-mediated actin filament elongation at the pointed ends instead of the barbed end). Strikingly, we observed that myosin concentration was enhanced, but the overall actin network contraction was significantly reduced compared to the architecture previously simulated and observed *in vivo* (Fig. 6D-F). Importantly, the forces acting on formin-elongated actin filaments were severely reduced (Fig. 6G). Indeed, with this network architecture, myosin motors merely moved toward the pulse center, in a polarity sorting mechanism (Fig. 6D, (Wollrab et al. 2018)), rather than pulling actin filaments to generate forces. These simulations therefore showed that the rapid assembly—by RhoA-driven pulses of formins—of a polar network architecture drives efficient actomyosin network contractility, supporting the remodeling of cell shape during pulsed contractions.

Altogether, these simulations results show that actomyosin network architecture –largely governed by the kinetics of formin-mediated actin filament assembly— controls the mechanics of pulsed contraction, thereby playing a key role to support the cellular function of pulsed contractions.

## Discussion

Precise architectural organization of the actomyosin network is crucial for force generation at the cell cortex. How such architectures are assembled in a dynamic network with fast turnover stands as a multiple answer question during development where force deployment is critical for embryo morphogenesis. Here, we show that formins organize a polar actin network during cortical pulsed contraction, in a biochemical system primed for rapid assembly.

Our results are based on a detailed description of the kinetics of actin assembly by formins. We show that formins elongate actin filaments at 1.2 μm·s^-1^, or ~450 monomers·s^-1^. Formin-mediated actin filament elongation *in vitro* has been proposed to overcome diffusion limiting rates (Drenckhahn & Pollard 1986), likely by allowing formins to “explore” a larger volume to “find” monomers (Courtemanche 2018). Assuming that elongation rates scale with the concentration of actin (in our experimental configuration formins are not anchored and unlikely to be directly subject to mechanical forces), in a solvent-independent manner (i.e. independently of viscosity—affecting the diffusion rate, or crowding effects, where solutes in the cell do not affect elongation rates of CYK-1), then formin velocity may provide a good indicator of the modulations of free G-actin concentration in the cell. Recent work however has shown that under saturating conditions, at concentrations of actin > 200 μM formins velocity could actually prove robust to variations in G-Actin and profilin concentration (Funk et al. 2019)).

Our analyses further revealed that two distinct populations with specific mobilities are recruited at the cell surface: superdiffusive and subdiffusive formins. We attributed these populations to elongating and recruited formins populations, respectively. Upon binding with RhoA, cytoplasmic formins would bind to the cortex, diffuse locally, bind an available filament barbed-end and start elongation. Kinetically, therefore, we expected the observe the sequential recruitment of recruited **then** elongating formins, but instead observe the opposite sequence. To explore how this dynamics could emerge, we developed a biochemical model to see if we could reconcile our biochemical scheme with our observations. Our model showed that the two models come together under a specific set of assumptions, where barbed ends available for elongation are limiting, formins are recruited in large numbers and the conversion reaction is fast compared to other reactions in the system. And while our approach does not exclude other possible models—for example that formins are initially elongating, then somehow stall after some time— the set of assumptions we designed seems robust to variations in the biochemical activation scheme used. While this analysis of CYK-1 dynamics provides a new and interesting perspective on the dynamics of barbed ends at the cortex during activation by RhoA, we still lack tools to conclusively explore a collection of issues: when are barbed ends generated and by which mechanisms, what is the dynamics of capping during pulses, how many barbed ends are generated? We also do not have yet the resolution to explore the specific nature of the observed barbed ends: are formins capable of hetero-dimerizing or co-assembling with other factors (e.g. CapZ), and could this lead to the formation of inactive barbed ends, these other formins acting as competitive inhibitors for CYK-1 formin at the filament barbed end? Our results however clearly show that the actin in *C. elegans* is biochemically primed for rapid response. When the RhoA signaling cascade is activated, actin assembly is saturated by formins to drive an efficient and optimally rapid response to signaling cues.

Strikingly, the geometry of the assembled network is controlled by the geometry of the upstream signaling factors: local RhoA activation drives the assembly of a polar actin network. This suggests that, at this scale, actin architecture seems to be fundamentally driven by the spatial patterning of assembly kinetics, rather than by a reorganization of the actin cytoskeleton by the mechanical motor activity of Myosin II (Reymann et al. 2016).

The architecture of the assembled structure is well tuned for actomyosin contractility, with Myosin II recruited at the center of the prospective pulsed contraction (Fig. S2), while actin is assembled in a polar aster network, with barbed ends pointing outwards of the assembled architecture. Our numerical simulations show that, while actomyosin networks can generate tension in the absence of a specific network architecture, actomyosin networks perform differently depending on their organization and the contractile efficiency of actomyosin network remains functionally linked to their geometry. Therefore, the actin assembly transduction machinery downstream of RhoA converts a chemical RhoA gradient into a polar actin network architecture, with a structure well adapted to the contractile function of actomyosin pulses in morphogenesis.

With precisely timed cell cycles, similar in duration to the ones in Drosophila syncytial embryo, lasting ~10’, *C. elegans* embryonic early development cell cycles unfold very fast (Brauchle et al. 2003)—compared to other early embryos, for instance in mouse embryos early cell cycles last about 20 h (Yamagata & FitzHarris 2013), sea urchin 150 min (Chassé et al. 2016) or even ascidians (~30 min) (Dumollard et al. 2013). In *C. elegans*, the 10 min cycles are divide roughly equally, into ~5 min for mitosis and 5 min interphase with cortical pulses. As a consequence, cell polarity, compartmentalization, and cell shape changes are heavily constrained in time. An actin network primed for fast assembly, together with the polar architecture of actomyosin pulsed contractions, may set the stage for rapid and efficient contractions and cell shape changes. During gastrulation, this very same organization may thus drive a fast apical constriction, and a subsequent timely internalization of endodermal cells.

Actomyosin network contractility is a key conserved feature of eukaryotic cells. Biochemically, the contractile structure assembled in *C. elegans* is very similar to the nodes assembled in fission yeast during contractile ring assembly: formin actin-filament elongators, Myosin II motors and actin cables (Vavylonis et al. 2008; Munro et al. 2004). However, several key differences separate the two contractile modules. Structurally, the size of the biological systems diverge strongly. At the level of the cell, a fission yeast cell spans ~ 14 μm long and 3 μm wide during cell division, against 50 μm in length and 30 μm in width for the *C. elegans* embryo (Fig 7A,B). The two contractile macromolecular assemblies are also very different: fission yeast nodes are <600nm wide and initially distant by <1 μm on average, while *C. elegans* actomyosin pulses are 3-5 μm wide and separated by 5-10μm (Fig. S2, (Michaux et al. 2018; Naganathan et al. 2014)). The two systems are also biochemically distinct. The fission yeast formin Cdc12p elongates actin filaments with high processivity (koff ~7.10^-5^ s^-1^) but slow speed (10.6 monomers·s^-1^ at 1.5 μM [Actin], 4 μM [SpPRF], fission yeast profilin), while the *C. elegans* formin CYK-1 elongates actin filaments with a lower processivity (koff ~4.10^-3^ s^-1^) but much higher speed (63.2 monomers·s^-1^ at 1.5 μM [Actin] and 4 μM [PFN-1], the *C. elegans* profilin). In the *Search-Pull-Capture-Release* model, actin filament elongation takes place from a static barbed end (Pollard & Wu 2010; Vavylonis et al. 2008). However, in *C. elegans*, a similar mechanism would result in filament buckling or stalling in actin filament elongation. To drive the same functional output— contraction—the molecular homologs assemble a structurally distinct, geometrically opposite, architecture which is tuned to the scale of the biological system, revealing here an interesting instance of the tinkering of evolution.

**Figure 7.**
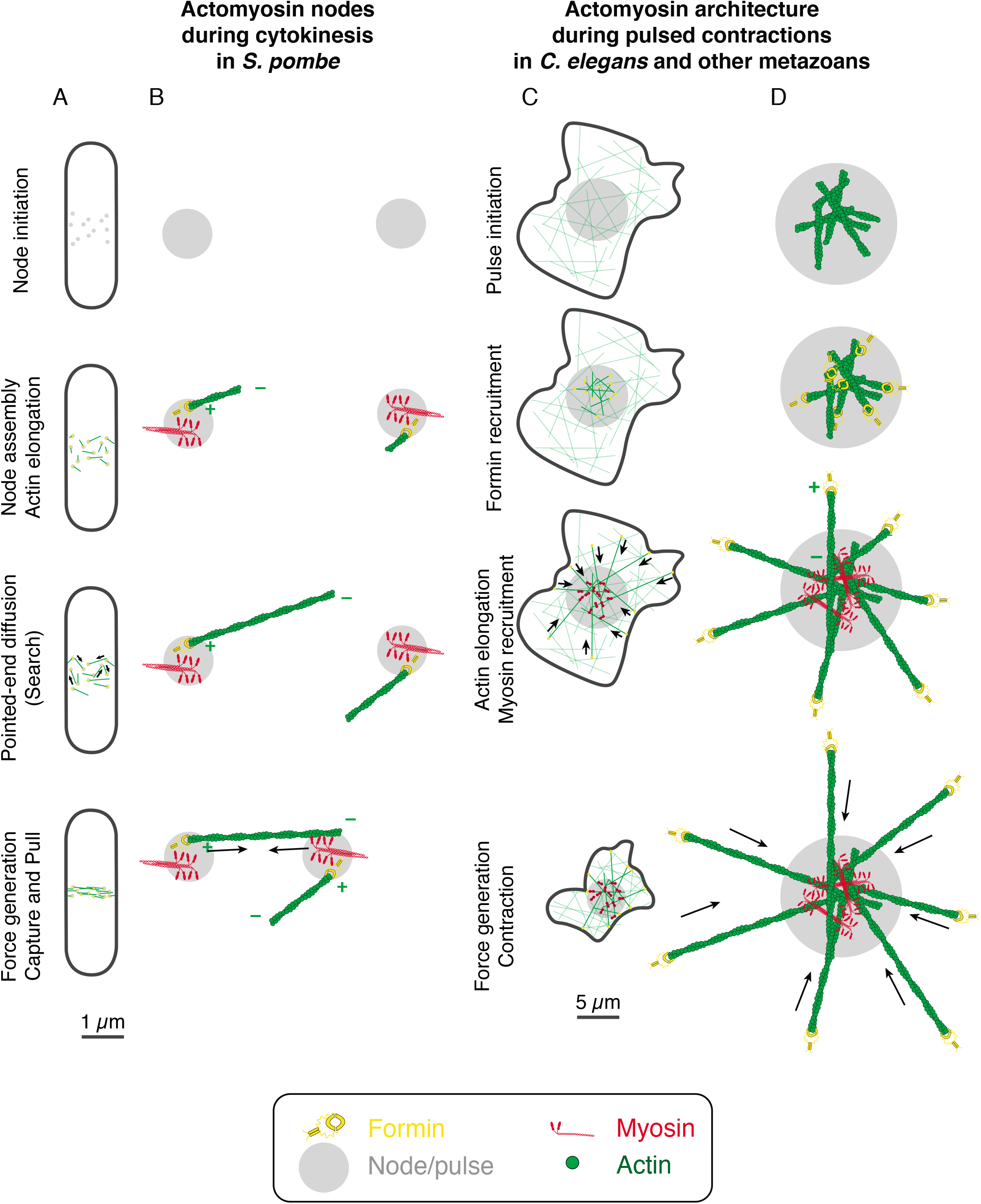
Compared analysis of contractility between pulsed contractility in metazoans (*Drosophila, C. elegans*) and yeast. **(A)** During cytokinesis in *S. pombe*, nodes form, cluster and align, forming the contractile to drive cell division, with a process scale size of 1-2 μm. Comparatively, in metazoan, pulsed contractions drive apical constriction over 10 to 50 μm micrometers. **(B)** In *S. pombe*, the formin Cdc12p is recruited in the nodes and drives filament elongation. Pointed-ends of filaments are proposed to explore until they are captured and pulled by Myo2p myosin filaments of another node. **(C)** During pulsed contractions in *C. elegans*, Rho recruits formins, which elongate actin filaments, followed by Myosin recruitment in the pulse center. Processive actin elongation by formins recruited at the pulse by Rho drives the formation of a polar actin network initiated at the pulse and extending over >10 μm from the pulse, with actin barbed-end pointing outwards. Myosin recruited by Rho in the center of the pulse then efficiently drives actin network contraction, pulling on actin cables assembled during the pulse to “reel in” the network towards the center of the pulse.

## Acknowledgements

We thank Anne-Cecile Reymann, Stephan Grill, Ronen Zaidel-Bar and Geraldine Seydoux for worm strains. We thank Marie Breau and Michel Labouesse for critical reading of the manuscript, Michel Labouesse, Marie Breau and members of the Labouesse laboratory for valuable discussions, as well as members of the Robin laboratory for discussions and technical assistance. We thank the IBPS imaging facility, and in particular Susanne Bolte, and France Lam, for excellent technical support.

This work used the Extreme Science and Engineering Discovery Environment (XSEDE, (Towns et al. 2014)), which is supported by National Science Foundation grant number ACI-1548562. The computations were conducted on the Comet supercomputer, which is supported by NSF award number ACI-1341698, at the San Diego Supercomputing Center (SDSC).

This work was supported by the ATIP/Avenir, ERC-Tremplin programs (to F.B. Robin) and Marie Curie Program MSCA-IF (661451).

The authors declare no competing financial interests.

## Material and Methods

### *C. elegans* culture and strains

See attached Supplementary Table 3.

### RNA interference

We performed RNAi using the feeding method as previously described (Timmons & Fire 1998). Bacteria targeting *perm-1* and *gfp* were obtained from the Kamath feeding library (Kamath et al. 2003).

The L4417 plasmid targeting *perm-1* and the entire GFP sequence (generated by the Fire lab and available at http://www.addgene.org/1649/) were transformed into HT115(DE3) bacteria. Bacterial cultures for feeding were grown for 10–12 h and then induced on standard nematode nutritional growth media plates containing 50 μg/ml ampicillin and 1 mM IPTG for 16–24 h at 20–25 °C, then stored at 4 °C. For *perm-1* RNAi, L4 stage larvae were placed on feeding plates for 16–24 h before imaging.

### Imaging conditions

We dissected gravid hermaphrodites and mounted one-cell embryos under #1.5 22-mm square coverslips in 2.5 μl of water or standard Egg Salts buffer (118 mM NaCl, 40 mM KCl, 3.4 mM CaCl2, 3.4 mM MgCl2, 5 mM HEPES, pH 7.4) containing ~500 uniformly sized polystyrene beads (15.6 ± 0.03 μm diameter, Bangs labs, #NT29N) to achieve uniform compression of the embryo surface across experiments (Robin et al. 2014).

We performed near-TIRF imaging at 19–21°C on an inverted Nikon Ti-E N-Storm microscope, equipped with motorized TIRF illuminator, Apo TIRF 100x Oil-immersion DIC N2 objective (Nikon) with 1.49–numerical aperture (NA), and PFS-S Perfect Focus unit (Nikon). Laser illumination at 488 nm and 561 nm from 300 mW solid-state sapphire laser (Coherent) was set at 30 % of maximal power and delivered by fiber optics to the TIRF illuminator. Images were magnified by a 1.5× lens and collected on an Andor iXon Ultra DU-897 EMCCD camera, yielding a pixel size of 107 nm.

We controlled laser illumination angle and intensity and image acquisition using NIS Elements software (Nikon). For all experiments, we set the laser illumination angle to a standard value that was chosen empirically to approximately maximize signal intensity while maintaining even illumination across the field of view. For all SPT experiments, we collected images in streaming mode with continuous illumination at 15-60% laser intensity (100% ≈ 1.6 μW·μm^-2^) with 50 ms exposures to achieve frame rates of 20 frames/s.

### Tuning GFP levels to achieve single-molecule densities

The quasi-steady-state densities observed during imaging depend on the initial (unobserved) densities, photobleaching rates and the intrinsic exchange kinetics of the target molecule (see main text and below). We thus determined the appropriate initial densities empirically for a given strain and experiment. We achieved these initial densities by using two methods as previously described (Robin et al. 2014). For SWG282 (CYK-1::GFP over-expression), we used RNAi directed against the GFP sequence to deplete the pool of GFP-tagged proteins. RNAi against maternal proteins typically yields an exponential decrease in the maternal protein with time of exposure (Oegema & Hyman 2006). We controlled the degree of depletion by synchronizing larvae and sampling embryos at different times after the initiation of feeding to identify times at which discrete diffraction-limited speckles were observed at the cell surface. The optimal time was relatively consistent across experiments for a given strain and varied from 12–36 h depending on transgene expression levels and relative abundance at the cell surface vs. cytoplasm. To fine-tune density levels, we used brief (<10 s) pulses in epi-illumination mode at high laser power until adequate density was reached (Robin et al. 2014).

### Drug perfusion experiments

For exposing embryos to 10 μM of Latrunculin A (Sigma L5163) or to 10 μg/mL of Nocodazole (Sigma M1404) in Egg Salts buffer during image acquisition, we used wider coverslips (22 mm x 30 mm) so that a perfusion chamber is formed between coverslip and slide, and the coverslip passes about 3 mm from the side of the slide. On the inverted microscope, this outer side of coverslip helps as support to able to deposit the perfusion volume as a drop (4 μL) while imaging. The drug solution likewise perfused by capillarity between slide and coverslip exposes the embryos to the drug instantly. The perfusion timepoint is visible in the corresponding movies as a brief brightfield illumination and used as a reference during analysis.

### Assessing potential adversary effects of compression, laser exposure and GFP fusion

We followed experimental procedures as previously tested (Robin et al. 2014). Using photobleaching to reduce GFP-tagged protein levels from full to single-molecule levels in one step resulted in arrested development. However, the laser exposure required to fine-tune densities by photobleaching, or that occurring during single-molecule imaging, did not cause embryos to arrest. In all of our single-molecule imaging experiments, we verified that embryos initiated and completed cytokinesis with normal timing or, in the case of nocodazole treated embryos multiple nuclei were present in the cell. To confirm that no adverse effects on population dynamics were associated with GFP fusion in the CYK-1::GFP CRISPR strain, we also used a CYK-1::mNeon CRISPR fusion to confirm our results (data not shown, strain available upon request).

### Single-molecule detection and tracking

We used a publicly available Matlab implementation of the Crocker-Grier algorithm for single-particle detection and tracking (Pelletier et al. 2009; Crocker & Grier 1996). In brief, the Crocker-Grier method localizes particles to subpixel resolution in individual frames by fitting local intensity peaks to a Gaussian point spread function. The two key detection parameters—peak and mean intensity of the candidate particles—are adjusted empirically for given imaging conditions using a graphical user interface. The particles are then linked frame to frame by minimizing the global displacement across all particles, given a user-chosen cutoff value for maximum particle displacement. A second parameter, the gap size, allows the possibility of ignoring ‘gaps’ in a trajectory due to transient failures to detect particles. These transient failures occur mainly because motion blur causes the particle intensity to fall transiently below the detection threshold.

To estimate actin concentration, we assumed that elongation rates scale linearly with actin concentration, and used previously measured elongation rates of 60 monomers·s^-1^, at 1.5 μM ATP-Actin (Neidt, Scott, et al. 2008; Neidt, Skau, et al. 2008).

To infer filament length, we made the following assumptions:

1. actin monomers display simple mono-exponential half-life at the cortex,
2. actin monomers display a half-life measured by tracking and smPReSS of 0.08-0.15 s (Robin et al. 2014; Michaux et al. 2018),
3. elongating formins display simple mono-exponential half-life at the filament barbed end,
4. elongating formins display a half-life measured by smPReSS of ~0.11 s (Fig. 2D).

Under these assumptions, we consider solely actin filaments assembled by formins. From the perspective of an actin monomer at the time of disassembly, two options are possible:

1. the monomer disassembles while the formin is still elongating,
2. the formin unbinds and elongation stops before the monomer disassembles.

The “effective” elongation time of the formin on the filament is then the minimum value between (1) and (2). If actin lifetime and formin mediated actin-filament elongation time have independent exponential distributions of parameters 1/τ_actin_ and 1/τ_formn_, then the minimum between the two values also has exponential distribution of parameter 1/τ_filament_ = 1/τ_actin_ +1/τ_formin_. Under these conditions, the length of the elongated filament is:

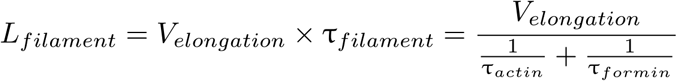

### Formin Speed measurement analysis

We performed single-molecule imaging as described previously. We mounted the embryos between glass slides with squares wells of 20 μm thick Epoxy and #1.5 coverslips (170 μm thick) in 2.5 μL of 0.22 μm filtered water with 15.4 μm polystyrene beads. We imaged single molecules using 50% of 90mW of 488 nm laser, 50 ms of exposure, no delay between frames, using Photometrics 95B prime 22 mm sCMOS camera. Laser angle was set to 65°. Room temperature maintained between 19 and 20.5 °C. After acquisition, we averaged two consecutive frames, in order to achieve 10 frames per second using ImageJ software (NIH Image, Bethesda, MD). We used Matlab implementation of the Crocker-Grier algorithm (Crocker & Grier 1996) by the Kilfoil lab for single-particle tracking. We selected manually a ROI to exclude tracks from residual particles outside the cell for each stage: whole embryo for one-cell stage, anterior (AB) cell for two-cell stage, posterior AB daughter cell (ABp) for four-cell stage. Each stage was separated into three phases based on observed cortical dynamics: interphase (pulsed contractions at the cortex); mitosis (cortex “stable” with no identifiable pulsed contractions); and cytokinesis (visualized by cleavage furrow assembly).

Subsequent image analysis was performed in Matlab. We selected the trajectories based on their anomalous diffusion coefficient *D* and scaling exponents α. Tracks were classified as subdiffusive and superdiffusive, and selected specifically superdiffusive trajectories. In order to calculate the velocity only during elongation of actin filaments, we performed a second selection to exclude tracks displaying multiple behaviors during their lifetime (due to switches between subdiffusive and superdiffusive) and retained tracks displaying exclusively superdiffusive behavior. Finally, we screened individual trajectories manually to retain tracks that were closer to a line to avoid skewing our estimates of particle speed.

### Statistical analysis

23 to 40 tracks per embryo were selected. Normal distributions were verified. Two-sample Student tests (t-tests) were performed to measure the significance of the difference in speed between each stage and phase. *** means p<0.001, ** means p<0.01, ns: non-significant.

### Two-color imaging microscopy

We performed single-molecule imaging as described previously. Acquisitions were performed with the Andor iXon 897 EMCCD camera. We imaged at 30% of 90 mW for 488 nm and 561 nm, with 50 ms exposure and no delay between frames (100 ms between two successive frames of the same channel). After acquisition, we averaged five consecutive frames, in order to achieve 2 frames per second using ImageJ. Pulsed contractions were selected manually and data from the intensity profile of a line drawn through the pulse is collected in a single frame. 3 pulses per embryo in 6 embryos (total of 18) were analyzed in Matlab (R2018a version). Intensities are smoothened and normalized with maximum being 1 and immediately preceding minimum being 0. Data for myosin (NMY-2::mKate2, in red channel) is aligned at 0,98% of the maximum and this alignment is propagated to the corresponding data in the green channel.

### Tracking of individual pulses of CYK-1::GFP

We used a semi-automatic approach to identify and follow CYK-1 pulses during the two-cell stage interphase, in the anterior blastomere. We manually identified isolated pulses and drew a ROI over the surface of each pulse (about 6 μm in diameter), at about 5 frames before maximum contraction of the area can be detected. The ROI was then automatically propagated in time before and after t0, and also we designed a Matlab script (A-STAR Methods PipelineSimPulse) allowing to adapt automatically the surface of the ROI in order to include the full trajectory of particles appearing within the ROI. To eliminate drift of the ROI associated with cortical flows, independently of CYK-1 mobility within the pulse area, we used a dedrifting routine on each particle based on the displacement of its neighbors. This was important for mobility analysis as particles registered with a global drift—and therefore displaying a persistent directional motion—would otherwise register as superdiffusive.

### Single particles tracking and pulse analysis pipeline in Matlab

We designed an analysis pipeline based on Matlab scripts (A-STAR Methods PipelineSimPulse, code available upon request) that includes CYK-1::GFP particles detection and tracking, reduction to the surface of the embryo and the AB cell, de-drifting of the trajectories and MSD analysis for segregation in different mobility populations (mainly superdiffusive CYK-1 vs. subdiffusive CYK-1). The further step is to intersect the matrix of all these trajectories with the specific ROI of each pulse. The final step is to normalize and align all the pulses (number of particles in time) with respect to their maximum and the minimum number of particles before, then to measure the angle orientation of every vector formed by the trajectories with respect to the center of the ROI.

### Numerical simulation of formin local recruitment

We used MATLAB to compute a 2D simulation of local formin activation and actin filament elongation. Pulses were spatially and temporally distributed in an embryo’s shaped mask in a random manner. Pulse were defined by a fixed 5×5 μm window (100×100 pixels) and a 20 seconds time window (400 frames). Pulses could not overlap in time and space. Using experimental data, density of formin recruitment, position around the pulse center and kinetics of recruitment could be computed in each pulse. We added 0.01 formins recruitment / μm^2^/frame all over the embryo mask (independent of pulses generation) corresponding to the formin recruitment rate observed in areas away from a pulse. According to experimental data, 80% of the formin recruited were assigned to be subdiffusive while 20% of them were assigned superdiffusive. Since we aimed to study superdiffusive particles, we approximate sub-diffusive and diffusive particles as a unique population of immobile particles. Each position of superdiffusive tracks were computed using the following sequence. The length of the step Rn was picked in a normal distribution whose mean is 1.23 μm/s and standard deviation is 0.30 μm/s (values extracted from experimental data).

The orientation θ_n_ of the step was calculated assuming a persistent length P_L_ of 15μm (close to the actin persistence length, (Howard 2001)).

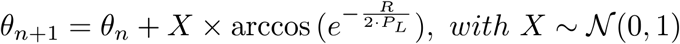

Length of track were assigned using the distribution of track length for sub-diffusive and super-diffusive particles, respectively. Simulated data were analyzed using the same methods as experimental data.

### Overview of the computational model of actomyosin mechanics

For simulations in this study, we used a well-established agent-based model of actomyosin networks based on the Langevin equation (J. Li et al. 2017; Jung et al. 2015; T. Kim et al. 2009; Mak et al. 2016). The detailed descriptions about the model and all parameters used in the model are explained in Supplementary Text and Table S1. In the model, actin filament (F-actin), motor, and ACP are coarse-grained using cylindrical segments (Fig. S4A). The motions of all the cylindrical segments are governed by the Langevin equation for Brownian dynamics. Deterministic forces in the Langevin equation include bending and extensional forces that maintain equilibrium angles formed by segments and the equilibrium lengths of segments, respectively, as well as a repulsive force acting between neighboring pairs of segments for considering volume-exclusion effects.

The formation of F-actin is initiated by a nucleation event, followed by polymerization at the barbed end and depolymerization at the pointed end. ACPs bind to F-actin without preference for cross-linking angles at a constant rate and also unbind from F-actin at a force-dependent rate determined by Bell’s law (Bell 1978). Each arm of motors binds to F-actin at a constant rate, and it then walks toward the barbed end of F-actin or unbinds from F-actin at force-dependent rates determined by the parallel cluster model (Erdmann et al. 2013; Erdmann & Schwarz 2012). For all simulations in this study, we used a thin computational domain (20×20×0.1 μm) with periodic boundary conditions only in x and y directions (Fig. S4B). In z direction, the boundaries of the domain exert repulsive forces on elements that moved beyond the boundaries. At the beginning of each simulation, a thin actin network is formed via self-assembly of F-actin and ACP.

For implementing RhoA activation, the domain is divided into 16 subdomains (4×4 in x and y directions). Every 30 s, one of the subdomains is randomly selected and then activated. In the activated subdomain, a fraction of the barbed ends of F-actins are randomly chosen and then undergo faster polymerization by a factor, *ϱ*_f_, for the duration of *τ*_f_. With the reference values of *ϱ*_f_ = 10 and *τ*_f_ = 10 s, F-actins are elongated by ~10 μ_M_ on average. After the time delay of *d*_M_, motors in the activated subdomain are allowed to self-assemble into thick filament structures for the duration of *τ*_M_. The reference values of *d*_M_ and *τ*_M_ are 5 s and 15 s, respectively. These active motors in the form of thick filaments can contract the part of the network in the activated subdomain. Once they become inactive after *τ*_M_, the motors are disassembled into monomers that cannot bind to F-actin.

## Bibliography

Agarwal, P. & Zaidel-Bar, R., 2019. Principles of Actomyosin Regulation In Vivo. Trends in Cell Biology, 29(2), pp.150–163.

Bell, G.I., 1978. Models for the specific adhesion of cells to cells. Science, 200(4342), pp.618–627.

Bement, W.M. et al., 2015. Activator-inhibitor coupling between Rho signalling and actin assembly makes the cell cortex an excitable medium. Nature Cell Biology, 17(11), pp.1471–1483.

Bement, W.M., Benink, H.A. & Dassow, von G., 2005. A microtubule-dependent zone of active RhoA during cleavage plane specification. The Journal of Cell Biology, 170(1), pp.91–101.

Benink, H.A., 2005. Concentric zones of active RhoA and Cdc42 around single cell wounds. The Journal of Cell Biology, 168(3), pp.429–439.

Bidone, T.C. et al., 2017. Morphological Transformation and Force Generation of Active Cytoskeletal Networks. A. R. Asthagiri, ed. PLoS computational biology, 13(1), p.e1005277.

Blanchoin, L. et al., 2014. Actin dynamics, architecture, and mechanics in cell motility. Physiological Reviews, 94(1), pp.235–263.

Brauchle, M., Baumer, K. & Gönczy, P., 2003. Differential activation of the DNA replication checkpoint contributes to asynchrony of cell division in C. elegans embryos. Current Biology, 13(10), pp.819–827.

Burkel, B.M. et al., 2012. A Rho GTPase signal treadmill backs a contractile array. Developmental Cell, 23(2), pp.384–396.

Carvalho, A. et al., 2011. Acute drug treatment in the early C. elegans embryo. B. Lehner, ed. PLoS ONE, 6(9), p.e24656.

Chassé, H. et al., 2016. Cyclin B Translation Depends on mTOR Activity after Fertilization in Sea Urchin Embryos. PLoS ONE, 11(3), p.e0150318.

Coravos, J.S. & Martin, A.C., 2016. Apical Sarcomere-like Actomyosin Contracts Nonmuscle Drosophila Epithelial Cells. Developmental Cell, 39(3), pp.346–358.

Courtemanche, N., 2018. Mechanisms of formin-mediated actin assembly and dynamics. Biophysical reviews, 10(6), pp.1553–1569.

Courtemanche, N. et al., 2013. Tension modulates actin filament polymerization mediated by formin and profilin. Proceedings of the National Academy of Sciences of the United States of America, 110(24), pp.9752–9757.

Crocker, J.C. & Grier, D.G., 1996. Methods of digital video microscopy for colloidal studies. Journal of Colloid and Interface Science, 179(1), pp.298–310.

Drenckhahn, D. & Pollard, T.D., 1986. Elongation of actin filaments is a diffusion-limited reaction at the barbed end and is accelerated by inert macromolecules. The Journal of biological chemistry, 261(27), pp.12754–12758.

Dumollard, R. et al., 2013. Beta-catenin patterns the cell cycle during maternal-to-zygotic transition in urochordate embryos. Developmental Biology, 384(2), pp.331–342.

Erdmann, T. & Schwarz, U.S., 2012. Stochastic force generation by small ensembles of myosin II motors. Physical Review Letters, 108(18), p.188101.

Erdmann, T., Albert, P.J. & Schwarz, U.S., 2013. Stochastic dynamics of small ensembles of non-processive molecular motors: the parallel cluster model. The Journal of Chemical Physics, 139(17), p.175104.

Funk, J. et al., 2019. Profilin and formin constitute a pacemaker system for robust actin filament growth. eLife, 8, p.1826.

Galkin, V.E. et al., 2011. Remodeling of actin filaments by ADF/cofilin proteins. Proceedings of the National Academy of Sciences of the United States of America, 108(51), pp.20568–20572.

Galkin, V.E. et al., 2010. Structural polymorphism in Factin. Nature Structural & Molecular Biology, 17(11), pp.1318–1323.

He, L. et al., 2010. Tissue elongation requires oscillating contractions of a basal actomyosin network. Nature Cell Biology, 12(12), pp.1133–1142.

Higashida, C., 2004. Actin Polymerization-Driven Molecular Movement of mDia1 in Living Cells. Science, 303(5666), pp.2007–2010.

Higgs, H.N., 2005. Formin proteins: a domain-based approach. Trends in biochemical sciences, 30(6), pp.342–353.

Howard, J., 2001. Mechanics of motor proteins and the cytoskeleton, Sinauer Ed, pp. 110–111.

Jégou, A., Carlier, M.-F. & Romet-Lemonne, G., 2013. Formin mDia1 senses and generates mechanical forces on actin filaments. Nature Communications, 4, p.1883.

Jung, W. et al., 2015. F-actin cross-linking enhances the stability of force generation in disordered actomyosin networks. Comp. Part. Mech., 2, pp.317–327.

Kamath, R.S. et al., 2003. Systematic functional analysis of the Caenorhabditis elegans genome using RNAi. Nature, 421(6920), pp.231–237.

Kim, H.Y. & Davidson, L.A., 2011. Punctuated actin contractions during convergent extension and their permissive regulation by the non-canonical Wnt-signaling pathway. Journal of Cell Science, 124(4), pp.635–646.

Kim, T., 2015. Determinants of contractile forces generated in disorganized actomyosin bundles. Biomechanics and Modeling in Mechanobiology, 14(2), pp.345–355.

Kim, T. et al., 2009. Computational analysis of viscoelastic properties of crosslinked actin networks. D. A. Fletcher, ed. PLoS computational biology, 5(7), p.e1000439.

Koenderink, G.H. & Paluch, E.K., 2018. Architecture shapes contractility in actomyosin networks. Current Opinion in Cell Biology, 50, pp.79–85.

Kreten, F.H. et al., 2018. Active bundles of polar and bipolar filaments. Physical Review E, 98(1-1), p.012413.

Kubota, H. et al., 2017. Biphasic Effect of Profilin Impacts the Formin mDia1 Force-Sensing Mechanism in Actin Polymerization. Biophysj, 113(2), pp.461–471.

Lecuit, T. & Lenne, P.-F., 2007. Cell surface mechanics and the control of cell shape, tissue patterns and morphogenesis. Nature Reviews Molecular Cell Biology, 8(8), pp.633–644.

Lenz, M., Gardel, M.L. & Dinner, A.R., 2012. Requirements for contractility in disordered cytoskeletal bundles. New Journal of Physics, 14(3), p.033037.

Lenz, M., Thoresen, T., et al., 2012. Contractile Units in Disordered Actomyosin Bundles Arise from FActin Buckling. Physical Review Letters, 108(23), p.238107.

Li, F. & Higgs, H.N., 2005. Dissecting requirements for auto-inhibition of actin nucleation by the formin, mDia1. The Journal of biological chemistry, 280(8), pp.6986–6992.

Li, F. & Higgs, H.N., 2003. The mouse Formin mDia1 is a potent actin nucleation factor regulated by autoinhibition. Current Biology, 13(15), pp.1335–1340.

Li, J. et al., 2017. Buckling-induced F-actin fragmentation modulates the contraction of active cytoskeletal networks. Soft Matter, 13(17), pp.3213–3220.

Linsmeier, I. et al., 2016. Disordered actomyosin networks are sufficient to produce cooperative and telescopic contractility. Nature Communications, 7, p.12615.

Maddox, A.S. et al., 2005. Distinct roles for two C. elegans anillins in the gonad and early embryo. Development, 132(12), pp.2837–2848.

Maître, J.-L. et al., 2015. Pulsatile cell-autonomous contractility drives compaction in the mouse embryo. Nature Cell Biology, 17(7), pp.849–855.

Mak, M. et al., 2016. Interplay of active processes modulates tension and drives phase transition in self-renewing, motor-driven cytoskeletal networks. Nature Communications, 7, p.10323.

Martin, A.C., Kaschube, M. & Wieschaus, E.F., 2009. Pulsed contractions of an actin-myosin network drive apical constriction. Nature, 457(7228), pp.495–499.

Mayer, M. et al., 2010. Anisotropies in cortical tension reveal the physical basis of polarizing cortical flows. Nature, 467(7315), pp.1–7.

Mi-Mi, L. et al., 2012. Z-line formins promote contractile lattice growth and maintenance in striated muscles of C. elegans. The Journal of Cell Biology, 198(1), pp.87–102.

Michaux, J.B. et al., 2018. Excitable RhoA dynamics drive pulsed contractions in the early C. elegansembryo. The Journal of Cell Biology, 217(12), pp.4230–4252.

Miller, A.L. & Bement, W.M., 2009. Regulation of cytokinesis by Rho GTPase flux. Nature Cell Biology, 11(1), pp.71–77.

Munjal, A. & Lecuit, T., 2014. Actomyosin networks and tissue morphogenesis. Development, 141(9), pp.1789–1793.

Munro, E., Nance, J. & Priess, J.R., 2004. Cortical flows powered by asymmetrical contraction transport PAR proteins to establish and maintain anterior-posterior polarity in the early C. elegans embryo. Developmental Cell, 7(3), pp.413–424.

Naganathan, S.R. et al., 2014. Active torque generation by the actomyosin cell cortex drives left-right symmetry breaking. J. Ferrell, ed. eLife, 3, p.e04165.

Naganathan, S.R. et al., 2018. Morphogenetic degeneracies in the actomyosin cortex. eLife, 7, p.354.

Nance, J., 2003. C. elegans PAR-3 and PAR-6 are required for apicobasal asymmetries associated with cell adhesion and gastrulation. Development, 130(22), pp.5339–5350.

Nance, J. & Priess, J.R., 2002. Cell polarity and gastrulation in C. elegans. Development, 129(2), pp.387–397.

Neidt, E.M., Scott, B.J. & Kovar, D.R., 2008. Formin Differentially Utilizes Profilin Isoforms to Rapidly Assemble Actin Filaments. The Journal of biological chemistry, 284(1), pp.673–684.

Neidt, E.M., Skau, C.T. & Kovar, D.R., 2008. The cytokinesis formins from the nematode worm and fission yeast differentially mediate actin filament assembly. The Journal of biological chemistry, 283(35), pp.2387223883.

Nishikawa, M. et al., 2017. Controlling contractile instabilities in the actomyosin cortex. eLife, 6, p.058101.

Oegema, K. & Hyman, A.A., 2006. Cell division. WormBook, pp.1–40.

Olson, S.K. et al., 2012. Hierarchical assembly of the eggshell and permeability barrier in C. elegans. The Journal of Cell Biology, 198(4), pp.731–748.

Pelletier, V. et al., 2009. Microrheology of Microtubule Solutions and Actin-Microtubule Composite Networks. Physical Review Letters, 102(18), p.188303.

Piekny, A., Werner, M. & Glotzer, M., 2005. Cytokinesis: welcome to the Rho zone. Trends in Cell Biology, 15(12), pp.651–658.

Pollard, T.D. & Wu, J.-Q., 2010. Understanding cytokinesis: lessons from fission yeast. Nature Reviews Molecular Cell Biology, 11(2), pp. 149–155.

Pruyne, D. et al., 2002. Role of Formins in Actin Assembly: Nucleation and Barbed-End Association. Science, 297(5581), pp.612–615.

Rauzi, M., Lenne, P.-F. & Lecuit, T., 2010. Planar polarized actomyosin contractile flows control epithelial junction remodelling. Nature, 468(7327), pp.1110–1114.

Reymann, A.-C. et al., 2016. Cortical flow aligns actin filaments to form a furrow. eLife, 5, p.e17807.

Robin, F.B. et al., 2014. Single-molecule analysis of cell surface dynamics in Caenorhabditis elegans embryos. Nature Methods, 11(6), pp.677–682.

Roh-Johnson, M. et al., 2012. Triggering a cell shape change by exploiting preexisting actomyosin contractions. Science, 335(6073), pp.1232–1235.

Shekhar, S. et al., 2015. Formin and capping protein together embrace the actin filament in a ménage à trois. Nature Communications, 6(1), pp.1–12.

Swan, K.A. et al., 1998. cyk-1: a C. elegans FH gene required for a late step in embryonic cytokinesis. Journal of Cell Science, 111 (Pt 14), pp.2017–2027.

Timmons, L. & Fire, A., 1998. Specific interference by ingested dsRNA. Nature, 395(6705), pp.854–854.

Towns, J. et al., 2014. XSEDE: Accelerating Scientific Discovery, Computing in Science & Engineering, vol. 16, no. 5, pp. 62–74.

Vavylonis, D. et al., 2008. Assembly Mechanism of the Contractile Ring for Cytokinesis by Fission Yeast. Science, 319(5859), pp.97–100.

Wollrab, V. et al., 2018. Polarity sorting drives remodeling of actin-myosin networks. Journal of Cell Science, 132(4), p.jcs219717.

Yamagata, K. & FitzHarris, G., 2013. 4D imaging reveals a shift in chromosome segregation dynamics during mouse pre-implantation development. Cell cycle, 12(1), pp.157–165.

Yu, Q. et al., 2018. Balance between Force Generation and Relaxation Leads to Pulsed Contraction of Actomyosin Networks. Biophysical Journal, 115(10), pp.2003–2013.

