## Supplementary Information for "Rapid assembly of a polar network architecture drives efficient actomyosin contractility"

### Inventory of supplementary information

The following elements are provided as supporting information:

1. Figures S1–4, with legends,
2. Movies S1–10, with movie legend file,
3. Tables S1–3:
    a. Tables S1–2, statistical information pertaining to Figure 2,
    b. Table S3 summarizes all strains used in this study,
4. Method: Description of mathematical model of formin recruitment kinetics,
5. Method: Detailed description of the computational model of actomyosin mechanics,
6. Protocol: Flowchart of the image analysis procedure to study formin pulse dynamics.

### Supplemental Figures S1–S4

**Costache, Prigent Garcia et al.**

Figure S1

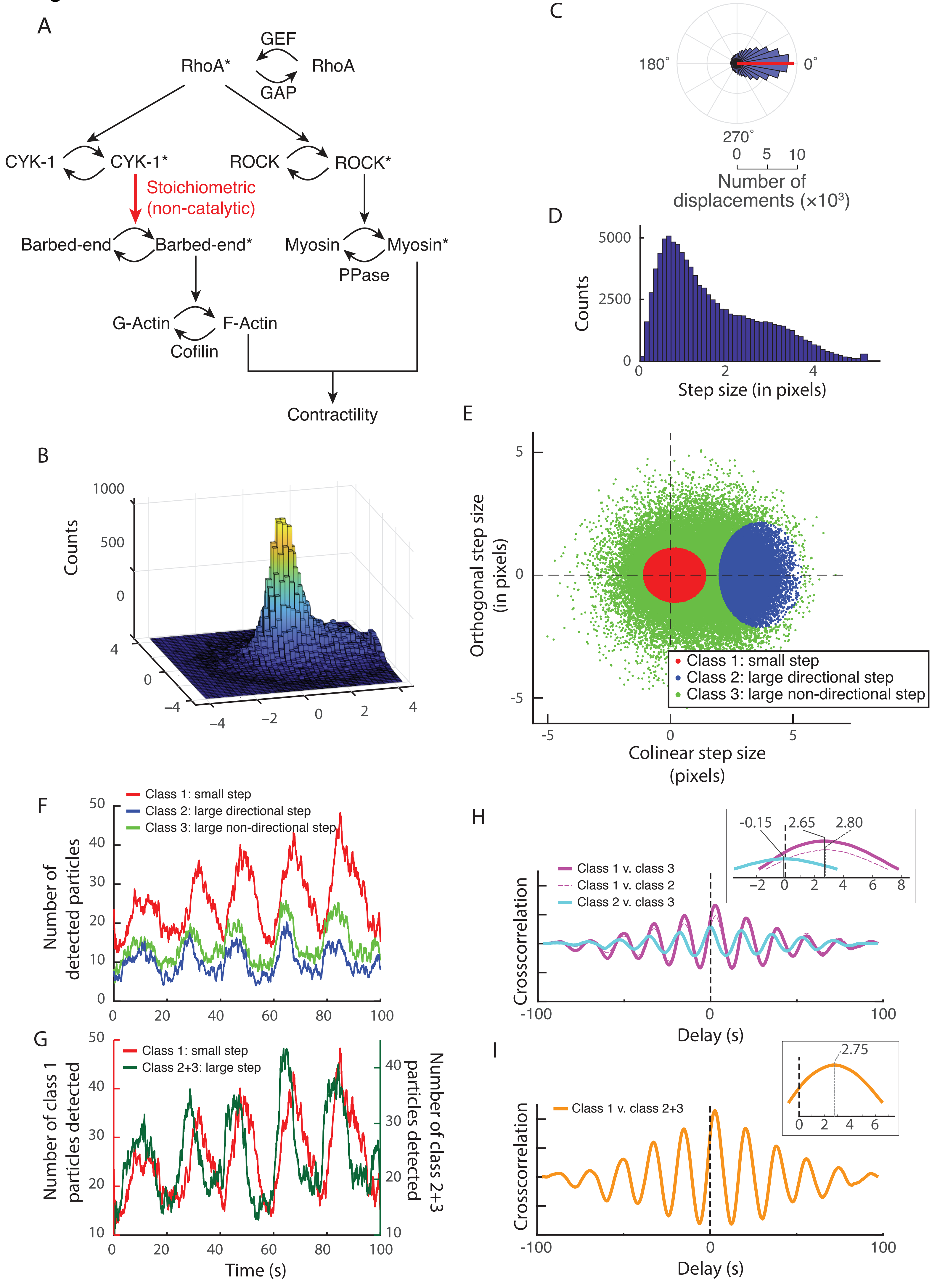
A
GEF
RhoA*
RhoA
GAP
CYK-1
CYK-1*
ROCK
ROCK*
Stoichiometric
(non-catalytic)
Barbed-end
Barbed-end*
Myosin
Myosin*
PPase
G-Actin
F-Actin
Cofilin
Contractility
B
Counts
1000
500
0
C
90˚
180˚
0˚
270˚
Number of displacements (×10³)
D
Counts
Step size (in pixels)
E
Orthogonal step size (in pixels)
Colinear step size (pixels)
Class 1: small step
Class 2: large directional step
Class 3: large non-directional step
F
Number of detected particles
G
Class 1: small step
Class 2+3: large step
Number of class 1 particles detected
Number of class 2+3 particles detected
Time (s)
H
Class 1 v. class 3
Class 1 v. class 2
Class 2 v. class 3
-0.15
2.65
2.80
Crosscorrelation
Delay (s)
I
Class 1 v. class 2+3
2.75
Crosscorrelation
Delay (s)

**Figure S1. Sorting and timing distinct populations of the formin CYK-1 reveals the kinetic signature of formin accumulation during pulsed contractions. (A)** Activation cascade of the formin CYK-1 by RhoA. **(B)** 3D histogram of step displacement with respect to the orientation of the previous step of the particle trajectory. Steps are taken of over 7 frames (350 ms). The histogram shows a large peak in zero, and a second additional peak around 3.5 pixels forward. **(C)** Distribution of angles of the steps in (B). **(D)** Distribution of step size, showing a merged but discernibly bimodal distribution of step sizes. **(E)** Automated classification of steps in (B) in 3 classes by 2-d gaussian fitting. Red: small/subdiffusive steps. Blue: large steps in the same direction as the previous step, corresponding to the second peak discussed in (B). Green: non-directional step, corresponding to formins changing course, or initiating elongation. **(F)** Temporal dynamics of the various step classes during a pulse, akin to Fig. 3A-C, but based on individual steps instead of whole trajectories. Green, red and blue as in (E). Note that Green and Blue curves, representing two classes of large steps (directional and non-directional) present similar temporal dynamics. **(G)** Temporal dynamics of subdiffusive steps vs "large steps"/superdiffusive steps. Dark green: merger of both classes of large steps (green and blue above). Red: small/subdiffusive steps. **(H, I)** Cross-correlation between the different populations of steps, and measured lags between these populations. Subdiffusive formins arrive ~2.7 s after superdiffusive (large steps) formins. Insets show an enlarged view of the peak region.

Figure S2

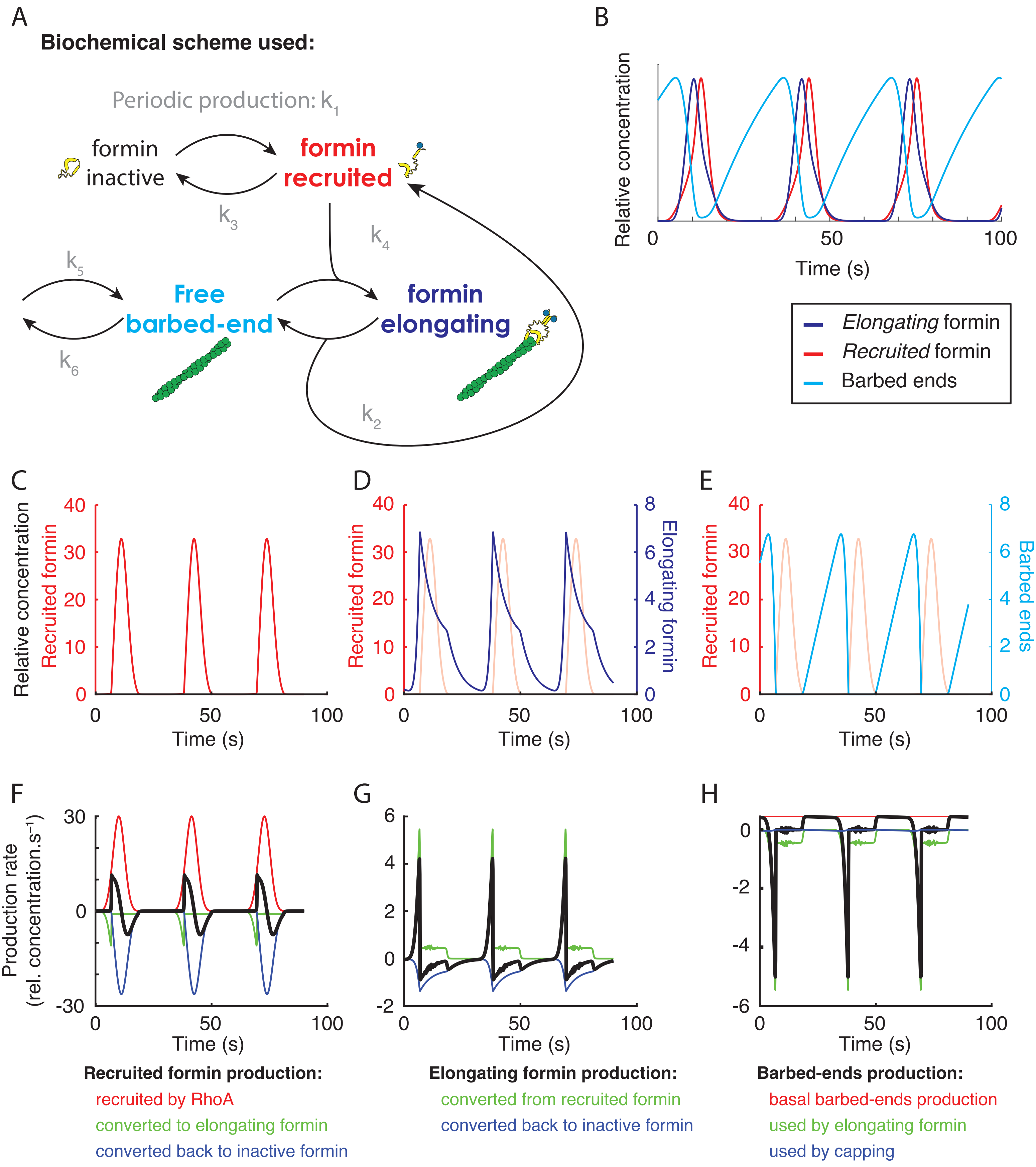

A
Biochemical scheme used:
Periodic production: k1
formin inactive
formin recruited
k3
k4
k5
Free barbed-end
formin elongating
k6
k2
B
Relative concentration
0
50
100
Time (s)
Elongating formin
Recruited formin
Barbed ends
C
Relative concentration
Recruited formin
40
30
20
10
0
50
100
Time (s)
D
Recruited formin
Elongating formin
8
6
4
2
Time (s)
E
Recruited formin
Barbed ends
Time (s)
F
Production rate
(rel. concentration.s-1)
30
0
-30
Time (s)
G
6
4
2
0
-2
Time (s)
H
0
-2
-4
-6
Time (s)
Recruited formin production:
recruited by RhoA
converted to elongating formin
converted back to inactive formin
Elongating formin production:
converted from recruited formin
converted back to inactive formin
Barbed-ends production:
basal barbed-ends production
used by elongating formin
used by capping

**Figure S2. Mathematical simulation of a barbed end depletion model – a detailed view. (A)** Biochemical scheme for formin recruitment, activation and formation of a complex with barbed ends. **(B)** An example of the evolution formin populations during a series of 3 pulses on the same graph. Cyan: Barbed ends, red: Recruited formins, purple: elongating formins. **(C-E)** Same as Fig. 4A-C, provided for reference for (F-H). **(C)** Temporal dynamics of recruited formins during a sequence of 3 pulses. Red: Recruited formins. **(D)** Same, with elongating formins (purple: elongating formins, light red: recruited formins). Recruited formins accumulate after elongating formins. **(E)** Temporal dynamics of barbed ends (Cyan: barbed ends, light red: recruited formins). **(F)** Sources and sinks affecting formin concentration are represented with a color code. Total derivative is presented in black, the positive contribution (production) and negative contribution (conversion) as described in the figure. **(G,H)** Same as (F) for the concentration of elongating formins (G) and barbed ends (H).

Figure S3

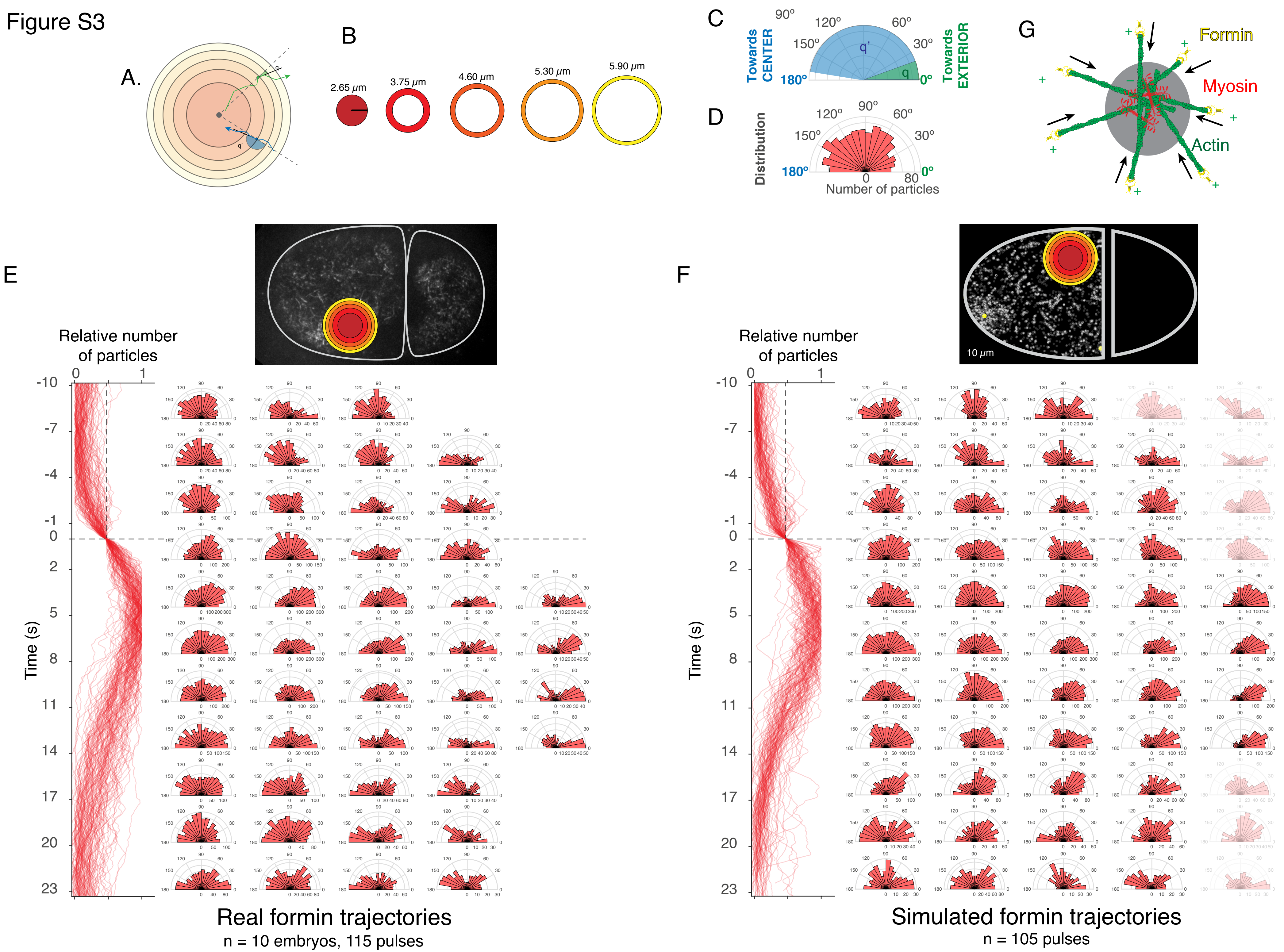

A.
B
2.65 μm
3.75 μm
4.60 μm
5.30 μm
5.90 μm
C
Towards CENTER
Towards EXTERIOR
90°
120°
60°
150°
30°
180°
0°
q'
q
D
Distribution
Number of particles
G
Formin
Myosin
Actin
E
F
Relative number of particles
Time (s)
10 μm
Real formin trajectories
n = 10 embryos, 115 pulses
Simulated formin trajectories
n = 105 pulses

**Figure S3**. **Simulated CYK-1 formin trajectories display similar polar orientation (barbed ends pointing out) during pulsed contractions, as measured in real embryos. (A)** Cartoon of a considered region of pulsed accumulation of formins, two examples of trajectories are drawn with the measured angle of the tangent from the center of the region to the vector of each particle in every position: green formin is directed outwards meanwhile blue formin is directed towards the center of the pulsed contraction region. **(B)** The pulsed contraction region is separated in concentric circular regions that have the same surface. The radius of each circular region is indicated. **(C)** Measure of the angle for two formin trajectories is performed with respect to the center of the pulse and the local orientation of the formin trajectory. The green track ($\theta \sim 25°$) is oriented with the barbed end of the filament pointing away from the center of the pulse, while the blue track ($\theta \sim 170°$) is oriented towards the center of the pulse. **(D)** All the measured angles of trajectories within a circular region from the pulsed contraction are displayed as distributions on 180° polar plots. **(E)** Measured angles of formins trajectories in pulsed contractions from real embryos, with respect to a spatial coordinate (horizontal) and a time coordinate (vertical) of the pulsed accumulation. The vertical red curves display the accumulation of formin particles for each pulse. All pulses are aligned at 45% ratio between minimum and maximum, at time 0, and we signal this time with a horizontal dashed line. Dataset is the same as Fig. 5. The regions around the pulse, and when pulse intensity is high (dashed black box), display a distribution skewed towards 0, showing that the orientation of the formins point outwards of the pulse region. **(F)** Similar analysis as in (E), with simulated formin tracks based on formin kinetics extracted from measurements in real embryos (see Methods section for details). **(G)** Mechanistic model of actin filament orientation during pulsed contractions.

Figure S4

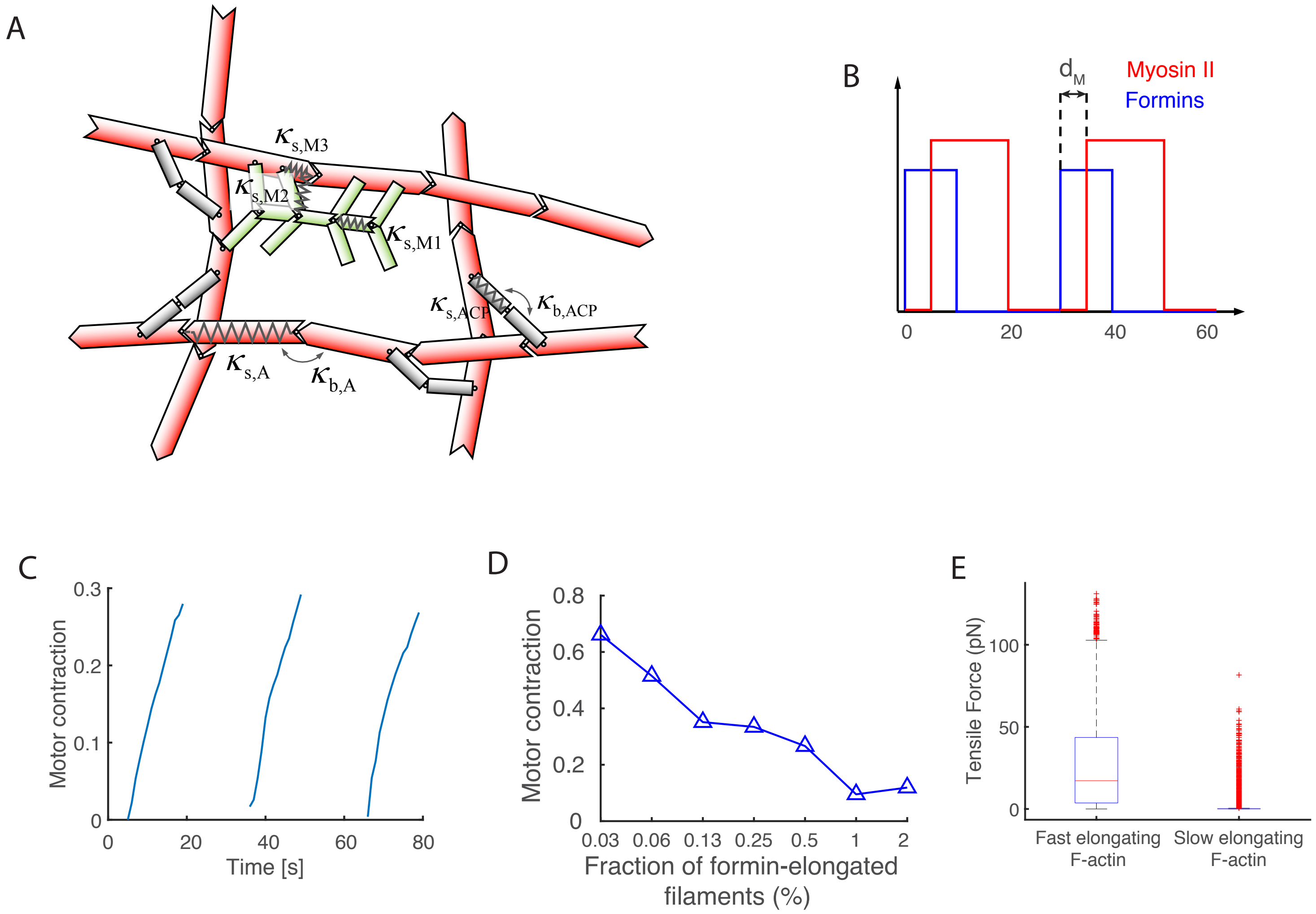

A
$\kappa_{s,M3}$
$\kappa_{s,M2}$
$\kappa_{s,M1}$
$\kappa_{s,ACP}$
$\kappa_{b,ACP}$
$\kappa_{s,A}$
$\kappa_{b,A}$
B
$d_M$
Myosin II
Formins
0
20
40
60
C
Motor contraction
0.3
0.2
0.1
0
0
20
40
60
80
Time [s]
D
Motor contraction
0.8
0.6
0.4
0.2
0
0.03
0.06
0.13
0.25
0.5
1
2
Fraction of formin-elongated filaments (%)
E
Tensile Force (pN)
100
50
0
Fast elongating F-actin
Slow elongating F-actin

**Figure S4. Numerical simulation of the mechanics of actomyosin networks during pulsed contractions. (A)** Agent-based model for simulating actomyosin networks. Actin (red), actin cross-linking protein (ACP), and motor (green) are simplified by cylindrical segments. Spring and bending forces with stiffness ($\kappa$s) maintain equilibrium lengths and angles, respectively. **(B)** Activation scheme of myosin and formins during pulsed contractions. **(C)** An example showing the extent of myosin contraction measured in three different activated regions. **(D)** Boxplot showing the distribution of tensile forces acting on fast elongating actin filaments (left) and slow elongating filaments (right) in the control condition. **(E)** The average of the maximum values of the motor contraction shown in (C) as a function of the fraction of quickly elongating filaments.

### Movies Legends

**Costache, Prigent Garcia et al.**

**Movies S1**. TIRF Single-molecule imaging of CYK-1::GFP formins localized at the cell cortex in the 2-cell stage *C. elegans* embryo. Different mobility behavior of the formins can be observed, some of them travel long distances in the cortex (white arrow), whereas some of them have a more confined mobility behavior (yellow arrow). Strain over-expresses CYK-1 fused with GFP. Scale bar is 5μm.

**Movie S2**. CYK-1::GFP formin trajectories can be easily observed by using a running time projection (bottom) of 100 time points (average of 4 frames, then running projection of 25 frames). Here is shown a 2-cell stage *C. elegans* embryo (TIRF imaging) during the pulsed contractions phase and the cortical stationary phase that follows. Strain over-expresses CYK-1 fused with GFP. Scale bar is 5μm.

**Movie S3**. Tracking of different mobility populations. Red circles indicate subdiffusive (confined) formins, green circles show the superdiffusive (active) formines and black circles indicate the formins that are present at the cortex for less than 15 frames in time and not categorized with a mobility profile. Strain over-expresses CYK-1 fused with GFP. Scale bar is 5μm.

**Movie S4**. Effect of drug treatment on formin mobility dynamics. TIRF microscopy of 2-cell stage *C. elegans* embryos expressing formin CYK-1::GFP (CRISPR) and permeabilized by *perm-1* RNAi. **(Left)** Embryo perfused with actin destabilizing drug Latrunculin A (10 μM) at indicated time (0min:45sec) on the movie. Following actin network destabilization, formins formed stable aggregates at the cell cortex, probably pulled by microtubules towards the cell center. **(Right)** Embryo perfused with microtubules destabilizing drug Nocodazole (10 $\mu g.ml^{-1}$) at indicated time (0min:45sec) on the movie. Formins still display normal mobility, after Nocodazole perfusion. In the projection of the same embryo (bottom), long formin trajectories are readily observed. Scale bar is 5μm.

**Movie S5**. Near-TIRF imaging of a 2-cell stage embryo during cortical stationary phase (mitosis) overexpressing actin fused with GFP. Acceleration x6. Scale bar: 10 μm. Actin molecules are rather immobile once recruited at the cortex.

**Movie S6**. Tracking of CYK-1 formins within a pulsed accumulation of formins in a 2-cell stage embryo using an adaptable region of interest. Green circles indicate subdiffusive, confined formins, whereas magenta circles show the superdiffusive, ballistic formins. Strain over-expresses CYK-1 fused with GFP.

**Movie S7**. Simulated 2-cell stage C. elegans embryo displaying CYK-1 formin trajectories as running projections. The tracks respect the spatial distribution (density, length, pulsed accumulations) as native formin trajectories, but the track orientation has been completely randomized. Yellow dots indicate formin pulsed accumulations.

**Movie S8**. A network without accelerated actin polymerization. In this movie, the magnitude of tensile forces acting on the network is visualized by color scaling; blue, white, and red represent low, intermediate, and high forces, respectively. Green represents active myosin motors. The duration of this movie is 80 s in the simulation.

**Movie S9**. A network with a moderate number of actin filaments undergoing fast polymerization. In this movie, the magnitude of tensile forces acting on the network is visualized by color scaling; blue, white, and red represent low, intermediate, and high forces, respectively. Active myosin motors are represented by green. The duration of this movie is 96 s in the simulation.

**Movie S10**. A network with a large number of actin filaments experiencing accelerated polymerization. In this movie, the magnitude of tensile forces acting on the network is visualized by color scaling; blue, white, and red represent low, intermediate, and high forces, respectively. Green indicates active myosin motors. The duration of this movie is 96 s in the simulation.

**Table S1: Statistical information supporting Fig. 2A,B, *Costache, Prigent-Garcia, et al.***

| | number of embryos | number of tracks | mean | median | high notch | low notch | std |
|---|---|---|---|---|---|---|---|
| 1c interphase | 6 | 240 | 1.2497 | 1.2641 | 1.2869 | 1.2412 | 0.1764 |
| 1c mitosis | 6 | 240 | 1.272 | 1.2896 | 1.3102 | 1.2689 | 0.1626 |
| 1c cytokinesis | 5 | 170 | 1.0496 | 1.0716 | 1.105 | 1.0382 | 0.2222 |
| 2c AB interphase | 7 | 280 | 1.2413 | 1.2385 | 1.2617 | 1.2153 | 0.1933 |
| 2c AB mitosis | 7 | 168 | 1.1848 | 1.1794 | 1.211 | 1.1479 | 0.1756 |
| 2c AB cytokinesis | 5 | 180 | 1.0442 | 1.026 | 1.0567 | 0.99534 | 0.1838 |
| 4c ABx interphase | 5 | 190 | 1.1546 | 1.155 | 1.1801 | 1.13 | 0.1675 |
| 4c ABx mitosis | 5 | 185 | 1.2407 | 1.2506 | 1.2764 | 1.2248 | 0.1768 |
| 2c P1 interphase | 7 | 161 | 1.1795 | 1.178 | 1.2086 | 1.1475 | 0.1849 |
| 2c P1 mitosis | 7 | 161 | 1.2176 | 1.2204 | 1.2519 | 1.1889 | 0.1929 |

**Table S2: Statistical information supporting Fig. 2A,B, *Costache, Prigent-Garcia, et al.***

| | paired | Two-sample ttest speed on tracks | Paired-sample ttest speed on mean | Two-sample ttest speed on mean |
|---|---|---|---|---|
| 1-cell interphase/ 1-cell mitosis | y | 0.1503 | 0.3676 | 0.5396 |
| 1c interphase/ 1c cytokinesis | | 9.92E-22 | | 0.0016 |
| 1c mitosis/ 1c cytokinesis | | 1.78E-27 | | 0.0013 |
| 2c AB interphase/ 2c AB mitosis | y | 0.0021 | 0.0082 | 0.3185 |
| 2c AB interphase/ 2c AB cytokinesis | | 1.12E-24 | | 0.0091 |
| 2c AB nitosis/ 2c AB cytokinesis | | 2.19E-12 | | 0.0429 |
| 4c ABp interphase/ 4c ABp mitosis | y | 1.91E-06 | 0.0328 | 0.0321 |
| 1c interphase/ 2c AB interphase | | 0.609 | | 0.8609 |
| 2c AB interphase/ 4c ABp interphase | | 6.99E-07 | | 0.0962 |
| 1c mitosis/ 2c AB mitosis | | 3.88E-07 | | 0.1 |
| 2c AB mitosis/ 4c ABp mitosis | | 0.0031 | | 0.3138 |
| 1c cytokinesis/ 2c AB cytokinesis | | 0.8027 | | 0.9341 |
| 2c P1 interphase/ 2c P1 mitosis | y | 0.071 | 0.4844 | 0.5465 |
| 2c AB interphase/ 2c P1 interphase | y | 0.0011 | 0.1845 | 0.2725 |
| 2c AB mitosis/ 2c P1 mitosis | y | 0.1066 | 0.1871 | 0.6054 |

**Table S3: List of strains used in *Costache, Pirgent-Garcia, et al.***

| Strain name | Genotype | Source |
|---|---|---|
| N2 | Wild-type Bristol strain | CGC |
| EM302 | *mgSi5[cb-UNC-119 (+) GFP::ANI-1(AH+PH)]II; nmy-2(cp52[nmy-2::mKate2 + unc-119(+)]) I; unc-119(ed3) III* | Michaux et al, 2018 |
| FBR104 | *cyk-1(jme06[cyk-1::mNeon])III* | This study |
| FBR106 | *cyk-1(jme06[cyk-1::mNeon])III; gesIs001[Pmex-5::Lifeact::mKate::nmy-2UTR, unc-119+]* | This study |
| FBR160 | *cyk-1(jme14[cyk-1::eGFP])III* | This study |
| FBR175 | *cyk-1(jme14[cyk-1::eGFP])III; nmy-2(cp52[nmy-2::mKate2 + unc-119(+)]) I; unc-119(ed3) III* | This study |
| JH1541 | *unc-119(ed4); pJH7.03 [unc-119; pie-1:GFP:actin::pie-1 3' UTR]* | Courtesy of G. Seydoux |
| LP229 | *nmy-2(cp52[nmy-2::mKate2 + LoxP unc-119(+) LoxP]) I; unc-119 (ed3) III* | Dickinson et al, 2017 |
| SWG001 | *gesIs001[Pmex-5::Lifeact::mKate::nmy-2UTR, unc-119+]* | Reyman et al, 2016 |
| SWG282 | *gesIs008[Pcyk-1::CYK-1::GFP::cyk-1UTR, unc-119+]* | This study |

**Methods**

### Description of mathematical model of formin recruitment kinetics

**Costache, Prigent Garcia et al.**

### Assembly of a polar network architecture downstream of a signaling cascade

Vlad Costache[1], Séréna Prigent-Garcia[1], Camille Plancke[1], Jing Li[2], Simon Bégnaud[1], Shashi Kumar Suman[1], Taeyoon Kim[2], and François B. ROBIN[1]

[1]CNRS UMR7622 and Inserm ERL 1156, Institut de Biologie Paris Seine (IBPS), Sorbonne Université, Paris, France.
[2]Weldon School of Biomedical Engineering, Purdue University, West Lafayette, Indiana.

January 1, 2021

#### 1 Assumptions

The proposed model is based on the following set of assumptions:

1. RhoA activation activity is represented as a smooth periodic function $(sin^6(\omega t)$,
2. inactive formins are activated by RhoA and recruited to the cortex, becoming "recruited",
3. CYK-1 formins are poor nucleators but good elongators – we considered formins do not efficiently nucleate new filaments under physiological conditions (in vitro actin assembly yields ∼1 new nucleated filament per 550 CYK-1 formin molecule at 2.5 $\mu$M actin and 2.5 $\mu$M profilin PFN-1, Neidt:2008df)
4. once recruited at the cortex, formins bind to barbed ends through a trimolecular reaction to drive actin assembly, becoming "elongating",
5. recruited formins unbind from the cortex to the cytoplasmic pool, (6) elongating formins unbind from the cortex to the cytoplasmic pool.

#### 2 Definitions

$[CYK-1_{recruited}] : [CYK1^*]$
$[CYK-1_{elongating}] : [CYK1^{**}]$
$[Barbedends] : [BE]$
$Period : T$

#### 3 Equations

$$\frac{d\left([CYK1^{*}]\right)}{dt} = k_1 \cdot \sin^6\left(\frac{\pi t}{T}\right) - k_3 \cdot [CYK1^{*}] - 2 \times k_4 \cdot [CYK1^{*}]^2 \cdot [BE] \quad (1)$$

$$\frac{d\left([CYK1^{**}]\right)}{dt} = k_4 \cdot [CYK1^{*}]^2 \cdot [BE] - k_2 \cdot [CYK1^{**}] \quad (2)$$

$$\frac{d\left([BE]\right)}{dt} = -k_4 \cdot [CYK1^{*}]^2 \cdot [BE] + k_5 - k_6 \cdot [BE] \quad (3)$$

#### 4 Origin of the terms

$$\frac{d\left([CYK1^{*}]\right)}{dt} = k_1 \cdot \sin^6\left(\frac{\pi t}{T}\right) - k_3 \cdot [CYK1^{*}] - 2 \times k_4 \cdot [CYK1^{*}]^2 \cdot [BE] \quad (4)$$

The **first term** describes the pulse activation by Rho, which is transient and has a period of ∼30s. The **second term** describes inactivation of the recruited species. The **third term** describes the conversion from recruited to elongating species, as a trimolecular reaction, corresponding to the assumption that cytoplasmic, inactive formins are monomeric and assemble as dimers upon binding with barbed-ends.

$$\frac{d\left([CYK1^{**}]\right)}{dt} = k_4 \cdot [CYK1^{*}]^2 \cdot [BE] - k_2 \cdot [CYK1^{**}] \quad (5)$$

The **first term** equates the conversion from inactive to active species (third term above), while the **second term** describes inactivation of the active species which is not converted back to recruited but is instead inactivated in the cytoplasmic species.

$$\frac{d\left([BE]\right)}{dt} = -k_4 \cdot [CYK1^{*}]^2 \cdot [BE] + k_5 - k_6 \cdot [BE] \quad (6)$$

The **first term** corresponds to the conversion of *free* barbed ends. These *free* barbed ends are likely to correspond to capped barbed ends: formins will displace the equilibrium and replace capping proteins, due to their higher affinity for barbed ends compared to capping proteins (*Neidt et al, 2008*). The **second term** corresponds to a low, continuous source/production of barbed ends. The **third term** corresponds to an inactivation of free barbed ends, representing the disappearance of barbed ends after some time. A typical value for that term will be on the order of magnitude of actin turnover rate ($\sim 0.1 - 1\ s^{-1}$)

### Methods
### Description of the computational model of actomyosin mechanics
**Costache, Prigent Garcia et al.**

#### SUPPLEMENTAL INFORMATION

##### Brownian dynamics via the Langevin equation

In our agent-based model, F-actin is simplified into serially connected cylindrical segments with barbed and pointed ends. Motors have a backbone structure with eight arms ($N_a = 8$) attached, and each of the motor arm represents four myosin heads. Therefore, the total number of myosin heads represented by one motor is 32, which is not quite different from 56 myosin heads in one non-muscle myosin thick filament [8]. The backbone and arms of the motors are also described by cylindrical segments. ACPs are comprised of two cylindrical arm segments.

The displacements of all the cylindrical segments are determined by the Langevin equation with the negligence of inertia:

$$\mathbf{F}_i - \zeta_i \frac{\mathrm{d}\mathbf{r}_i}{\mathrm{d}t} + \mathbf{F}_i^{\mathrm{T}} = 0 \qquad \text{(S1)}$$

where $\mathbf{r}_i$ is a position vector of the $i$th element, $\zeta_i$ is a drag coefficient, $t$ is time, $\mathbf{F}_i$ is a deterministic force, and $\mathbf{F}_i^{\mathrm{T}}$ is a stochastic force satisfying the fluctuation-dissipation theorem [9]:

$$\left\langle \mathbf{F}_i^{\mathrm{T}}(t)\mathbf{F}_j^{\mathrm{T}}(t) \right\rangle = \frac{2k_{\mathrm{B}}T\zeta_i\delta_{ij}}{\Delta t}\boldsymbol{\delta} \qquad \text{(S2)}$$

where $\boldsymbol{\delta}$ is a second-order tensor, $\delta_{ij}$ is the Kronecker delta, and $\Delta t = 1.15\text{x}10^{-5}$ s is a time step. The drag coefficients are calculated via an approximated form for cylindrical objects [10]:

$$\zeta_i = 3\pi\mu r_{c,i} \frac{3 + 2r_{0,i} / r_{c,i}}{5} \qquad \text{(S3)}$$

where $\mu$ is the viscosity of surrounding medium, and $r_{0,i}$ and $r_{c,i}$ are the length and diameter of segments, respectively. The positions of all the cylindrical segments are updated at each time step via the Euler integration scheme:

$$\mathbf{r}_i(t+\Delta t) = \mathbf{r}_i(t) + \frac{d\mathbf{r}_i}{dt}\Delta t = \mathbf{r}_i(t) + \frac{1}{\zeta_i}\left(\mathbf{F}_i + \mathbf{F}_i^{\mathrm{T}}\right)\Delta t \tag{S4}$$

**Deterministic forces**

Deterministic forces include extensional forces maintaining equilibrium lengths, bending forces maintaining equilibrium angles, and repulsive forces accounting for volume-exclusion effects between actin segments. The extensional and bending forces originate from the following potentials:

$$U_{\mathrm{s}} = \frac{1}{2}\kappa_{\mathrm{s}}(r - r_0)^2 \tag{S5}$$

$$U_{\mathrm{b}} = \frac{1}{2}\kappa_{\mathrm{b}}\left(\theta - \theta_0\right)^2 \tag{S6}$$

where $\kappa_s$ and $\kappa_b$ are extensional and bending stiffnesses, $r$ and $r_0$ is the instantaneous and equilibrium lengths of cylindrical segments, and $\theta$ and $\theta_0$ are instantaneous and equilibrium angles formed by segments. The equilibrium length of actin segments ($r_{0,A}$ = 140 nm) and an equilibrium angle formed by two adjacent actin segments ($\theta_{0,A}$ = 0 rad) are maintained by extensional ($\kappa_{s,A}$) and bending ($\kappa_{b,A}$) stiffnesses of actins, respectively. The reference value of $\kappa_{b,A}$ corresponds to the persistence length of ~9 μm [11]. The equilibrium length of ACP arms ($r_{0,ACP}$ = 23.5 nm) and an equilibrium angle formed by the two arm segments of each ACP ($\theta_{0,ACP}$ = 0 rad) are regulated by extensional ($\kappa_{s,ACP}$) and bending ($\kappa_{b,ACP}$) stiffnesses of ACPs, respectively. The equilibrium length of motor backbone segments ($r_{s,M1}$= 42 nm) and an equilibrium angle formed by adjacent backbone segments ($\theta_{0,M}$ = 0 rad) are maintained by extensional ($\kappa_{s,M1}$) and bending ($\kappa_{b,M}$) stiffnesses, respectively. The value of $\kappa_{s,M1}$ is equal to that of $\kappa_{s,A}$, whereas the value of $\kappa_{b,M}$ is larger than that of $\kappa_{b,A}$. The extension of each motor arm is regulated by the two-spring model with stiffnesses of transverse ($\kappa_{s,M2}$) and longitudinal ($\kappa_{s,M3}$) springs. The transverse spring maintains an equilibrium distance ($r_{0,M2}$ = 13.5 nm) between the endpoint of a motor backbone and an actin segment where the arm of the motor binds, whereas the longitudinal spring maintains a right angle between the motor arm and the actin segment ($r_{0,M3}$ = 0 nm).

The repulsive force is represented by a harmonic potential [3]:

$$U_r = \begin{cases} \frac{1}{2}\kappa_r \left(r_{12} - r_{c,A}\right)^2 & \text{if} \quad r_{12} < r_{c,A} \\ 0 & \text{if} \quad r_{12} \geq r_{c,A} \end{cases} \tag{S7}$$

where $\kappa_r$ is the strength of repulsive force, and $r_{12}$ is a minimum distance between two actin segments. Forces exerted on actin segments by bound motors and ACPs or by the repulsive force are distributed onto the barbed and pointed ends of the actin segments as described in our previous work [2].

#### Dynamics of ACPs

ACPs bind to binding sites located on actin segments every 7 nm without preference for cross-linking angles at a constant rate and also unbind from F-actin at a force-dependent rate determined by Bell's law [5]:

$$k_{\mathrm{u,ACP}} = \begin{cases} k^0_{\mathrm{u,ACP}} \exp\left( \dfrac{x_{\mathrm{u,ACP}} \left| \vec{F}_{\mathrm{s,ACP}} \right|}{k_{\mathrm{B}} T} \right) & \text{if } r \geq r_{0,\mathrm{ACP}} \\ k^0_{\mathrm{u,ACP}} & \text{if } r < r_{0,\mathrm{ACP}} \end{cases} \tag{S8}$$

where $\left|\vec{F}_{\mathrm{s,ACP}}\right|$ is a spring force acting on an ACP arm, $k^0_{\mathrm{u,ACP}}$ is the zero-force unbinding rate constant, $x_{\mathrm{u,ACP}}$ is sensitivity to an applied force, and $k_{\mathrm{B}}T$ is thermal energy. The values of $k^0_{\mathrm{u,ACP}}$ (= 0.115 $s^{-1}$) and $x_{\mathrm{u,ACP}}$ (= $1.04\times10^{-10}$ m) are determined based on filamin A [12].

**Dynamics of motors**

Motor arms bind to binding sites on actin segments at the rate of $40N_{\mathrm{h}}$ $s^{-1}$, where $N_{\mathrm{h}} = 8$ is the number of myosin heads represented by each motor arm. The walking ($k_{\mathrm{w,M}}$) and unbinding ($k_{\mathrm{u,M}}$) rates of the motor arms are determined by the parallel cluster model to mimic the mechanochemical cycle of non-muscle myosin II [6, 7]. The details of implementation and benchmarking of the parallel cluster model in our models are described in detail in our previous study [13]. Note that $k_{\mathrm{w,M}}$ and $k_{\mathrm{u,M}}$ are smaller with larger applied loads because motors exhibit a catch-bond behavior. The unloaded walking velocity and stall force of motors are ~140 nm/s and ~5.7 pN, respectively.

**Actin dynamics**

The formation of F-actin is initiated from a nucleation event with the appearance of one cylindrical segment with polarity (i.e., with barbed and pointed ends) in a random orientation perpendicular to the z direction. The polymerization and depolymerization of actins are simulated by the addition and removal of cylindrical segments, respectively, as in our previous studies [4]. The average length of F-actin ($<L_{f}>$) used in simulations is ~1 μm. This value is comparable to that estimated in our in vivo experiments. In addition, with the reference values of the rate constants for actin dynamics, each F-actin turns over every ~10 s.

**Contraction of actin**

In order to quantitatively analyze the network morphology, we evaluate the contraction of actin, using the spatial distribution of F-actins in activated regions whose dimension is 5×5 μm in x and y directions. First, the activated region is divided into $N_G \times N_G$ grids. We found that the optimal level of $N_G$ is 20. All grids are indicated by their own coordinate, (*i, j*). In each grid, we measure the intensity of actin segments, $\rho_A^{i,j}$. Then, the standard deviation of $\rho_A^{i,j}$ in all 400 grids is calculated and divided by the mean value of actin density, $\overline{\rho_A}$. The normalized value represents the extent of actin contraction:

$$\text{Actin contraction} = \frac{1}{\overline{\rho_A}} \sqrt{\frac{\sum_{i=1}^{N_G} (\rho_A^{i,j}\big|_{i,j=1...N_G} - \overline{\rho_A})^2}{N_G - 1}} \qquad \text{(Eq. S9)}$$

We calculate the time evolution of actin contraction by subtracting the initial value of actin contraction from the instantaneous value at each time step. From the time evolution curve, we obtain the maximal contraction and contraction at a plateau phase.

**Contraction of myosin motors**

We calculate the extent of motor contraction using the coordinates of the centers of motor thick filaments, $(\overline{x_{m,i}}, \overline{y_{m,i}})$. We calculate a distance between each thick filament and the center position of a currently activated region at each time step. We assume that the average of all the distances represents the rough size of motor clusters. The average is further divided by an initial value:

$$\text{Motor contraction} = \frac{r_i}{r_0} = \frac{\sum_{j=1}^{N_{m,j}} \sqrt{(x_{m,j} - \overline{x_{m,i}})^2 + (y_{m,j} - \overline{y_{m,i}})^2} \; / \; N_{m,j}}{\sum_{j=1}^{N_{m,0}} \sqrt{(x_{m,j} - \overline{x_{m,0}})^2 + (y_{m,j} - \overline{y_{m,0}})^2} \; / \; N_{m,0}} \qquad \text{(Eq. S10)}$$

In the time evolution of motor contraction, we average the maximum values of the motor contraction in all pulse periods to use it as an indicator for the extent of motor contraction.

**Table S1.** List of parameters employed in the model. For some of the parameters, references are provided if the parameters were determined based on specific previous studies.

| Symbol | Definition | Value |
|---|---|---|
| $r_{0,A}$ | Length of an actin segment | $1.4\times10^{-7}$ [m] |
| $r_{c,A}$ | Diameter of an actin segment | $7.0\times10^{-9}$ [m] [14] |
| $\theta_{0,A}$ | Bending angle formed by adjacent actin segments | 0 [rad] |
| $\kappa_{s,A}$ | Extensional stiffness of F-actin | $1.69\times10^{-2}$ [N/m] |
| $\kappa_{b,A}$ | Bending stiffness of F-actin | $2.64\times10^{-19}$ [N·m] [11] |
| $r_{0,ACP}$ | Length of an ACP arm | $2.35\times10^{-8}$ [m] [15] |
| $r_{c,ACP}$ | Diameter of an ACP arm | $1.0\times10^{-8}$ [m] |
| $\theta_{0,ACP}$ | Bending angle formed by two ACP arms | 0 [rad] |
| $\kappa_{s,ACP}$ | Extensional stiffness of ACP | $2.0\times10^{-3}$ [N/m] |
| $\kappa_{b,ACP}$ | Bending stiffness of ACP | $1.04\times10^{-19}$ [N·m] |
| $r_{0,M1}$ | Length of a motor backbone segment | $4.2\times10^{-8}$ [m] |
| $r_{0,M2}$ | Length of a motor arm | $1.35\times10^{-8}$ [m] |
| $r_{c,M}$ | Diameter of a motor arm | $1.0\times10^{-8}$ [m] |
| $\theta_{0,M}$ | Bending angle formed by motor backbone segments | 0 [rad] |
| $\kappa_{s,M1}$ | Extensional stiffness of a motor backbone | $1.69\times10^{-2}$ [N/m] |
| $\kappa_{s,M2}$ | Extensional stiffness 1 of a motor arm | $1.0\times10^{-3}$ [N/m] |
| $\kappa_{s,M3}$ | Extensional stiffness 2 of a motor arm | $1.0\times10^{-3}$ [N/m] |
| $\kappa_{b,M}$ | Bending stiffness of a motor backbone | $5.07\times10^{-18}$ [N·m] |
| $N_h$ | Number of heads represented by a motor arm | 4 |
| $N_a$ | Number of arms per motor | 4 |
| $k_{n,A}$ | Nucleation rate of actin | 0.001 [$\mu M^{-1}s^{-1}$] |
| $k_{p,A}$ | Polymerization rate of actin at the barbed end | 5 [$\mu M^{-1}s^{-1}$] |
| $k_{d,A}$ | Depolymerization rate of actin at the pointed end | 50 [$s^{-1}$] |
| $k^0_{u,ACP}$ | Zero-force unbinding rate constant of ACP | 0.115 [$s^{-1}$] [12] |
| $x_{u,ACP}$ | Sensitivity of ACP unbinding to applied force | $1.04\times10^{-10}$ [m] [12] |
| $\kappa_r$ | Strength of repulsive force | $1.69\times10^{-3}$ [N/m] |
| $\Delta t$ | Time step | $1.15\times10^{-5}$ [s] |
| $\mu$ | Viscosity of surrounding medium | $8.6\times10^{-1}$ [kg/m·s] |
| $k_BT$ | Thermal energy | $4.142\times10^{-21}$ [J] |
| $C_A$ | Actin concentration | 200 [μM] |
| $R_M$ | Ratio of motor concentration to $C_A$ | 0.01 |
| $R_{ACP}$ | Ratio of ACP concentration to $C_A$ | 0.04 |
| $<L_f>$ | Average length of F-actins | ~1 [μm] |
| $\rho_f$ | Enhancement factor for faster actin polymerization | 10 (5 in one simulation) |
| $\tau_f$ | Duration of faster actin polymerization | 10 s (5 s in one simulation) |
| $d_M$ | Time delay of myosin activation | 5 s (0 s in one simulation) |
| $\tau_M$ | Duration of myosin activation | 15 s |

1. Li, J., et al., *Buckling-induced F-actin fragmentation modulates the contraction of active cytoskeletal networks.* Soft Matter, 2017. **13**(17): p. 3213-20.
2. Jung, W., M. P Murrell, and T. Kim, *F-actin cross-linking enhances the stability of force generation in disordered actomyosin networks.* Comput Part Mech, 2015. **2**(4): p. 317-27.
3. Kim, T., et al., *Computational analysis of viscoelastic properties of crosslinked actin networks.* PLOS Comput Biol, 2009. **5**(7): p. e1000439.
4. Mak, M., et al., *Interplay of active processes modulates tension and drives phase transition in self-renewing, motor-driven cytoskeletal networks.* Nat Commun, 2016. **7**: p. 10323.
5. Bell, G.I., *Models for the specific adhesion of cells to cells.* Science, 1978. **200**(4342): p. 618-27.
6. Erdmann, T., P.J. Albert, and U.S. Schwarz, *Stochastic dynamics of small ensembles of non-processive molecular motors: the parallel cluster model.* J Chem Phys, 2013. **139**(17): p. 175104.
7. Erdmann, T. and U.S. Schwarz, *Stochastic force generation by small ensembles of myosin II motors.* Phys Rev Lett, 2012. **108**(18): p. 188101.
8. Tyska, M.J., et al., *Two heads of myosin are better than one for generating force and motion.* Proc Natl Acad Sci U S A, 1999. **96**(8): p. 4402-7.
9. Underhill, P.T. and P.S. Doyle, *On the coarse-graining of polymers into bead-spring chains.* J Non-Newtonian Fluid Mech, 2004. **122**(1): p. 3-31.
10. Clift, R., J.R. Grace, and M.E. Weber, *Bubbles, drops, and particles*. 2005: Courier Corporation.
11. Isambert, H., et al., *Flexibility of actin filaments derived from thermal fluctuations. Effect of bound nucleotide, phalloidin, and muscle regulatory proteins.* J Biol Chem, 1995. **270**(19): p. 11437-44.
12. Ferrer, J.M., et al., *Measuring molecular rupture forces between single actin filaments and actin-binding proteins.* Proc Natl Acad Sci U S A, 2008. **105**(27): p. 9221-6.
13. Kim, T., *Determinants of contractile forces generated in disorganized actomyosin bundles.* Biomech Model Mechanobiol, 2015. **14**(2): p. 345-55.
14. Kishino, A. and T. Yanagida, *Force measurements by micromanipulation of a single actin filament by glass needles.* Nature, 1988. **334**(6177): p. 74-6.
15. Meyer, R.K. and U. Aebi, *Bundling of actin filaments by alpha-actinin depends on its molecular length.* J Cell Biol, 1990. **110**(6): p. 2013-24.

**Protocol**

**Flowchart of image analysis procedure to study formin pulse dynamics**

**Costache, Prigent Garcia et al.**

### Flowchart of the image analysis procedure

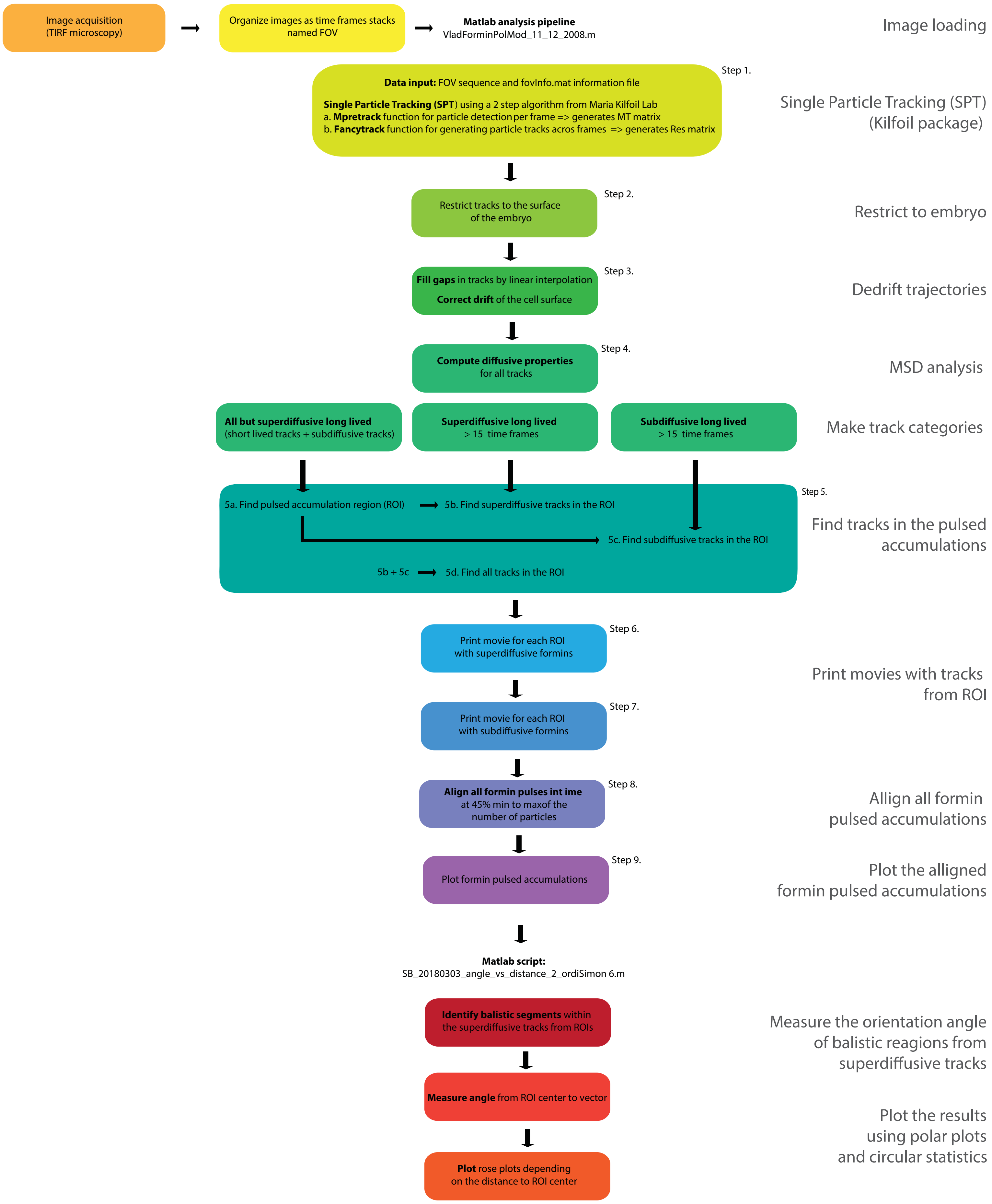

**Flowchart of the image analysis pipeline for measuring formin trajectories mobility and their orientation during pulsed accumulations at the cortex.**

All 9 steps are organized as code sections that include single particle tracking (SPT) using a previously published algorithm [1]. Data is further on managed as a matrix, throughout further processing steps: (1) elimination of particles outside the cell cortex, (2) correction of the drift and the MSD analysis (3) separation of the different mobility classes of formins : superdiffusive/ballistic (active formins), subdiffusive/confined (activated formins). Short-lived and subdiffusive particles serve as support to define the adaptive region of interest (ROI) corresponding to each pulsed accumulation region of formins. Pulses are then aligned in time with respect to the threshold ratio between minimum (before) and the maximum number of particles during the pulse, set at 45%. This analysis sets the stage for synchronizing across pulses the time sequences of arrival of the different formin populations at the cell cortex. A second script is used to measure the orientation angle between (1) the displacement vector of the active formins and (2) a vector starting at the center of the ROI/pulse and ending on the starting point of displacement. Results are printed as rose plot with circular statistics included. Circular statistics are performed using a matlab toolbox developed by Philipp Berens[2]. Images used in this analysis pipeline were obtained by near-TIRF microscopy at continuous 20 fps stream, mostly 16 bit, 6000 time frames stacks, 512x512 pixels, saved as single image sequences, named FOV that were directly recognized by Matlab during the first step of the pipeline.

1. V. Pelletier, N. Gal, P. Fournier, M. Kilfoil, *Phys. Rev. Lett.* **102**, 188303 (2009).
2. P. Berens, *Journal of Statistical Software*. **31**, 1–21 (2009).
